# Condenzymes: Biomolecular condensates with inherent catalytic activities

**DOI:** 10.1101/2024.07.06.602359

**Authors:** Michael W. Chen, Xiao Guo, Mina Farag, Naixin Qian, Xiaowei Song, Anton Ni, Vicky Liu, Xia Yu, Yuefeng Ma, Leshan Yang, Wen Yu, Matthew R. King, Joonho Lee, Richard N. Zare, Wei Min, Rohit V. Pappu, Yifan Dai

## Abstract

We report the discovery that chemical reactions can be catalyzed by condensates formed by intrinsically disordered proteins (IDPs). The proteins themselves lack any catalytic activities. Catalytic functions of condensates emerge as a consequence of sequence-dependent mesoscale electrochemical microenvironments created by phase separation. Stimulated Raman spectroscopy suggests that the catalytic behaviors of condensates are attributable to the spatial variations of water activities across condensate interiors and interfaces. We show that condensates are capable of catalyzing diverse cellularly relevant hydrolysis reactions. Through sequence design, the electrochemical properties of condensates can be programmed to exert control over catalytic behaviors. Incorporation of synthetic condensates into live cells alters transcription profiles and enables the activation of gene circuits that depend on products of hydrolysis reactions catalyzed by condensates. Our discovery of suggests that condensates, depending on their composition-dependent electrochemical properties, can be “Condenzymes”, which contribute unexpected emergent chemical functions in cells.

## INTRODUCTION

Biomolecular condensates concentrate condensate-specific biomolecules to regulate diverse cellular functions ^1^ such as stress responses and ribosomal biogenesis ^2–7^. Condensates form via composite processes that combine phase separation and reversible associations such as stoichiometric binding and non-stoichiometric percolation transitions ^8–10^. We refer to the collection of processes that contribute to condensate formation as condensation ^1,11,12^. Multivalent macromolecules, including intrinsically disordered proteins (IDPs) with specific molecular grammars ^6,7,11,13^, drive condensation.

Condensation, which involves spontaneous phase separation of the solution system as a whole, results in the coexistence of two or more phases ^12,14–21^. The differences in solvent and biomolecular compositions between the condensate interiors and coexisting dilute phases result in unique interphase physicochemical properties ^11,12,14,22,23^. These come about due to the establishment of phase equilibria whereby the chemical potentials of each species are equalized, thus giving rise to the possibility of asymmetric partitioning of these species in coexisting dense and dilute phases ^22^. The redistribution of solvent or asymmetry in net charge both lead to differential partition of solution ions across condensate interiors and coexisting dilute phases, which establish the interphase electric potentials such as Donnan potentials ^22–26^. These interphase potentials manifest as pH gradients, as shown for nucleoli and synaptic condensates, whereby the interiors are more acidic when compared to the surrounding nucleoplasm or dilute phase ^11,12,27^. Lastly, such electric potential gradients establish an electric double layer ^14^ at condensate surface where sharp transition of molecular density happens ^28^, and these have measurable electrochemical properties such as zeta potentials ^29^ and interfacial electric fields^14,30,31.^

The presence of unique interphase and interfacial properties, including asymmetries of ionic and hydration properties across coexisting phases and the presence interfacial electric fields^14,19–21,28^, suggests that distinct functionalities can be enabled by the unique chemical environments and electrochemical properties within condensates and their interfaces. This hypothesis derives from analogies with observations in colloid chemistry and micron-sized water droplets ^32–36^. A chemically active emulsion would imply that condensate functions are not solely determined by their capacity to store and compartmentalize biomolecules ^37,38^ or organize biochemical reactions. They also have the potential to contribute as drivers of biochemical reactions through their distinct physicochemical properties ^22^.

The growing inventory of distinct environmental features created within condensates and their interfaces inspired us to draw parallels with proteins enzymes ^23^, wherein the catalytic activities can be attributed to the local environments within the enzyme active sites ^39,40^. This includes the distinctive hydration environment and electric fields created by unique structural alignments of functional residues ^39–42^. We hypothesized that the physicochemical properties within condensate interiors and interfaces might share similarities to those within the catalytic sites of enzymes. Therefore, while enzymes are nanoscale catalysts, we conjectured that condensates have the potential to function as mesoscale catalysts. Accordingly, we tested whether the unique physicochemical features of condensates can enable spontaneous catalytic functions.

We present the results of assays and mass spectroscopy experiments used to evaluate the catalytic functions of different types of biomolecular condensates that possess distinct physicochemical features. The capability of condensates to catalyze hydrolysis reactions was evaluated both in vitro and in living cells. The underlying driving forces that mediate catalysis were studied using density functional theory and examined via vibrational Stark effect. Using *de novo* designs of IDPs that were guided by coarse-grained simulations, we uncover a decoupling of the contribution from the dense phase chemical environments and the interfacial electric fields to the kinetics of catalytic reactions driven by condensates. With stimulated Raman spectroscopy to map the local water hydrogen-bonding network, we found that the interior and the interface of condensates possess distinct water activity, which is sequence dependent. We further employed RNA sequencing, mass spectroscopy and synthetic gene circuits to demonstrate that the catalytic functions of condensates are capable of modulating downstream cellular processes. Overall, we find that condensates formed by catalytically inactive IDPs can drive the catalysis of biochemical reactions through their electrochemical properties. We name such condensates as “condenzymes”, biomolecular condensates with the inherent ability to catalyze chemical reactions.

## RESULTS

We first utilized two programmable model systems based on IDPs with repeating units namely, resilin-like polypeptides (RLPs) and elastin-like polypeptides (ELPs). RLPs undergo phase transitions driven by intra/inter-molecular interactions based on electrostatic and pi-based interactions ^43,44^, while ELPs undergo phase transitions driven by hydrophobicity based on the release of water molecules from the protein backbone ^45,46^. Therefore, the condensates formed by RLPs and ELPs arise from different driving forces, and they are defined by different chemical environments ^14,47^.

### Hydrolysis reactions can be catalyzed by condensates

Hydrolysis is a crucial biochemical process in live cells. For example, in the case of ATP hydrolysis, ATPases mediate the binding of ATP within the active site and cleavage of a bond by a water molecule ^48^. This enables energy production and is a key driver of cellular signaling ^49,50^. Many of the hydrolysis reactions favor an alkaline environment ^51^, and this matches the dense phase environments of RLP condensates^14^ (**Figure 1A**). Accordingly, we first explored the effects of condensates formed by the RLP_WT_ protein (amino acid sequence: Ser-Lys-Gly-Pro-[Gly-Arg-Gly-Asp-Ser-Pro-Tyr-Ser]_20_-Gly-Tyr) on ester hydrolysis. Phase transitions of RLP_WT_ were triggered by reducing the ionic activity of the solution (**Figure S1A, S1B**). We applied a chromogenic assay using *para*-nitrophenyl acetate (pNPA), which is a widely used substrate for the evaluation of esterase activity ^52^. Hydrolysis of pNPA decomposes the substrate to acetate and *para-*nitrophenol (pNP) (**Figure 1B**), and the product, pNP, can be directly monitored by UV-visible absorption spectroscopy at a wavelength of 410 nm, which is the absorption wavelength of phenol. We first verified the linear correlation between pNP concentration and absorbance at 410 nm (**Figure S1C**). Subsequently, we evaluated the pNPA hydrolysis reaction using an absorption scan under three different solution conditions: solution with condensates (*c*_RLP_ > *c*_sat_), solution without condensates (*c*_RLP_ < *c*_sat_), and the buffer solution without protein. Here, *c*_RLP_ and *c*_sat_ are the bulk and saturation concentrations of RLP, respectively. We confirmed the formation or the absence of condensates using differential confocal differential interference contrast microscopy (**Figure 1C**). In solutions with condensates, the colorless pNPA turned yellow after 120 min of incubation at room temperature (**Figure 1C**). This was evident from the appearance of a product absorption band at 410 nm (**Figure 1D**). In the solution without condensates and in the buffer solution, a similar intensity of the absorption was observed with a much lower signal compared to the solution with condensates. The weak signals in condensate-free or buffer solutions derive from the spontaneous hydrolysis of pNPA. The same trend was observed in time-dependent reaction tracking experiments (**Figure 1E**). Overall, our results suggest that the presence of condensates enhances the hydrolysis of pNPA.

**Figure 1.**
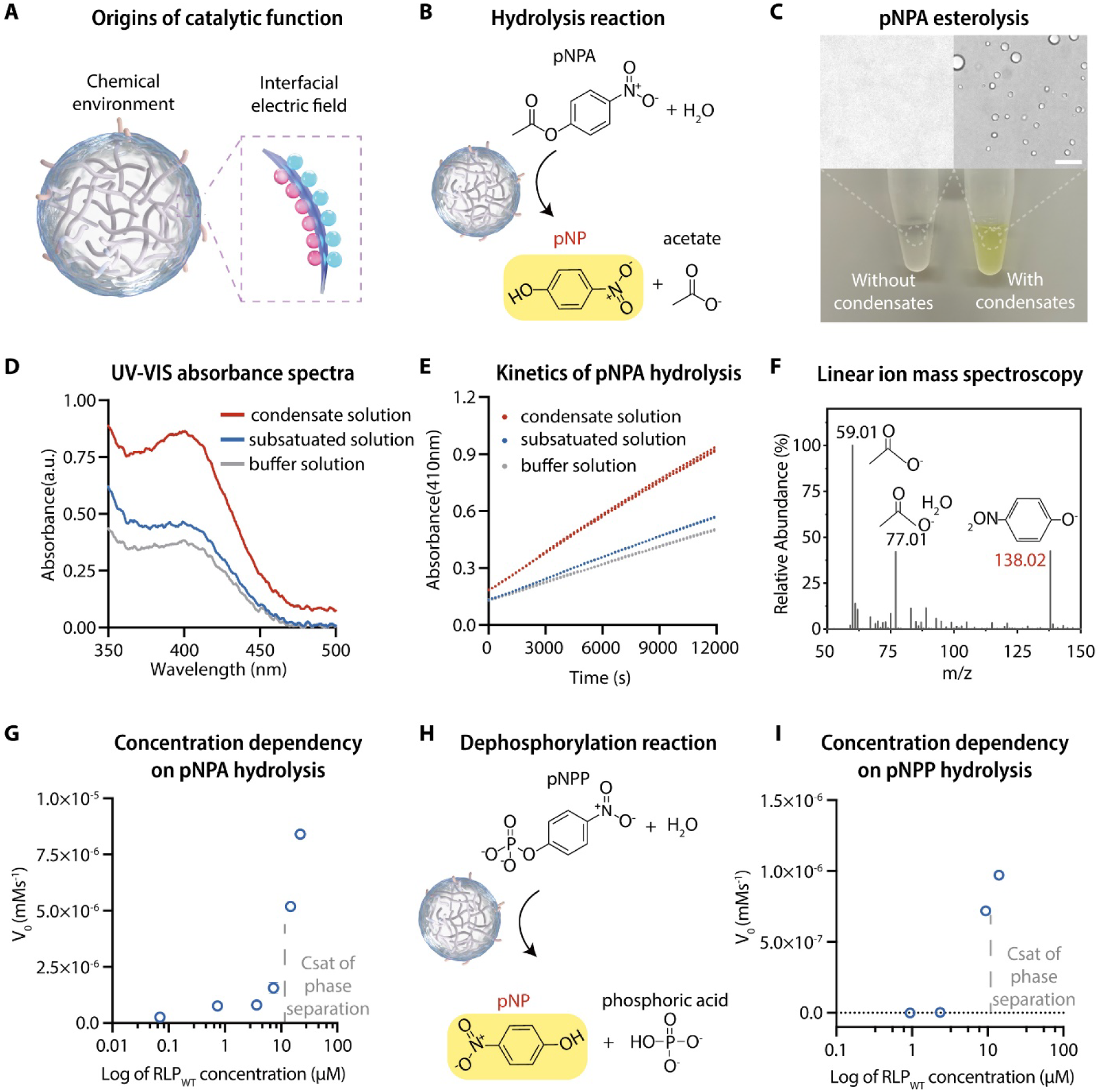
Biomolecular condensates can function as catalysts for hydrolysis reactions. **A,** Biomolecular condensates are defined by a unique chemical environment in the dense phase and an electric field at their interface. **B,** The hydrolysis reaction of *p-*nitrophenol acetate (pNPA) results in the formation of *p-*nitrophenol (a yellow product) and acetate. **C,** Image visualization of the catalytic reaction with a change of color in the solution with condensates. The samples containing 0.5 mM pNPA with or without condensates were incubated at room temperature for 120 min. The two images on the top are representative images displaying solution with or without condensates. Scale bar, 10 µm. **D,** UV-Vis absorbance spectrums of 1) the solution containing condensates and pNPA, 2) the solution containing only the dilute phase of a phase separated sample and pNPA, 3) the buffer solution containing pNPA. The final concentration of pNPA was 0.5 mM. **E,** Time-dependent absorbance tracking of pNPA hydrolysis with the same experimental setup as in d. N = 4 independent experimental trail. Each data point represents mean ± SD. **F,** A sample spectrum of linear ion mass spectroscopy analysis of the supernatant of the condensate solution containing pNPA after incubation of 120 min at room temperature. The same analysis was performed three times with different samples and the results were consistent. **G,** Protein concentration-dependent initial rate (V_0_) of the pNPA catalytic reaction. The protein concentration at which a sharp transition of the initial rate happened aligned with the saturation concentration of the RLP_WT_. **H,** The hydrolysis reaction of *p-*nitrophenol phosphate (pNPP) results in the formation of *p-* nitrophenol (a yellow product) and phosphoric acid. **I**, Protein concentration-dependent initial rate (V_0_) of the pNPP catalytic reaction.

To test if the reaction products are concordant with hydrolysis of pNPA, we used linear ion mass spectroscopy to analyze the supernatant of the condensate solution with pNPA after 120 min of incubation at room temperature. Distinct peaks at *m/z* ratios of 138, 77, and 59 were observed (**Figure 1F**). These correspond to *p*-nitrophenol, acetate, and acetate hydrate, confirming the esterolysis of pNPA in the solution with condensates. To evaluate whether the catalytic reaction is governed by condensate formation, we applied the pNPA assay to solutions containing different concentrations of RLP_WT_ and quantified the initial velocity (V_0_). The bulk concentration of RLP_WT_ is not linearly correlated with V_0_. Instead, we find that once the concentration of RLP_WT_ crosses its *c*_sat_ V_0_ increases exponentially (**Figure 1G**). This observation suggests that the catalytic function requires the formation of condensates.

To shed light on the mechanism by which condensates influence pNPA hydrolysis, we asked if the RLP_WT_ condensate acts as a catalyst instead of a reactant. As a catalyst, the material of the condensates should not be produced or destroyed by the pNPA decomposition reaction ^53^. We quantified the dense phase concentration using a sedimentation assay and compared the physical forms of the dense phase using confocal imaging. Across samples, with and without pNPA after 120 min of incubation at room temperature (**Figure S1D**), we did not observe differences in the dense phase concentrations or areas of the condensates (**Figure S2E, S2F**). These observations suggest that the reaction does not consume the components of the condensates and the constituents of the condensate are not chemically reactive to the substrate during the reaction period ^54^.

### Condensates can mediate hydrolysis of substrates with distinct side groups

To explore the capability of condensates on catalyzing hydrolysis reactions, we first investigated whether the RLP_WT_ condensates could mediate the hydrolysis of *p*-nitrophenyl phosphate (pNPP) (**Figure 1H**), which is a dephosphorylation process. Interestingly, we found that RLP_WT_ condensates can also catalyze dephosphorylation in a phase separation-dependent manner (**Figure 1I**). To further test the hydrolytic capability of condensates, we employed compounds with distinct chemical groups next to the ester bond as substrates (**Figure S1H**), including *p*-nitrophenyl butyrate (pNPB), *p*-nitrophenyl trimethylacetate (pNP-TMA) and *p*-nitrophenyl octanoate (pNPO). We found that at the same substrate concentration and reaction condition, the rate of hydrolysis reactions differ significantly between different substrates (**Figure S1H**). This observation suggests that condensate-substrate interaction might contribute to catalytic specificity (see Discussion). For those that could be hydrolyzed considerably by condensates, we found that the V_0_ of the hydrolysis of all the substrates increases exponentially once the RLP_WT_ concentration crosses the *c*_sat_ (**Figure S1I**), confirming that the hydrolytic behavior is a phase separation-dependent feature. These findings suggest that 1) condensates can drive different hydrolysis reactions and 2) the coupling of the chemical properties of the substrate and the condensate environment can be critical to the catalytic performance of condensates. This lends credence to the idea that the unique chemical features of condensates provide a favorable environment for enhancing the rates of reactions, similar to the functioning mechanism of an enzyme ^40^.

### Interfacial electric fields help lower the free energy barrier for hydrolysis reactions

The fact that condensates can catalyze different chemical reactions implies that the catalytic activities of condensates are influenced by the unique electrochemical features defined by the environments of condensate solution ^14^. We hypothesized that interfacial electric fields might one of the features of condensates that lower the activation free energy of reactions ^55,56^. To explore the physical basis of this possibility, we studied the influence of external electric field strengths on the rates of pNPA hydrolysis using Density Functional Theory (DFT) ^57^. We evaluated the difference in dipole moment between the initial substrate and the transition state of the pNPA to quantify the field-induced change of the reaction barrier (see STAR Methods for details). Under the influence of an electric field strength of ∼10^7^ V/cm (a typical value in the protein environment ^58^), the free energy barrier (ΔΔG^‡^) decreases from 9.26 kcal/mol to 7.73 kcal/mol at the intermediate state (**Figure S2**). This lowering of the barrier should correspond to an approximate 13-fold increase in the rate of pNPA hydrolysis, which is comparable to the observations in our experiments. This result suggests that the interfacial electric fields of condensates are likely to play an important role in catalyzing chemical reactions.

### The contributions of dense phase environment to hydrolysis

We set out to isolate the contributions of the microenvironment of the dense phase versus the interface to the catalytic functions of condensates. We first used the condensates formed by an ELP with a sequence of [Val-Pro-Gly-Val-Gly]_60_. Though the phase transition of ELP should generate a water density gradient, a limited pH gradient across the dilute and dense phases was observed due to the lack of charged residues ^14^. Compared to the RLP condensates, which possess a basic interior, ELP condensates possess a less favorable dense phase environment for alkaline hydrolysis reactions^23^. As expected, the ELP condensates were incapable of mediating the hydrolysis reaction (**Figure S3A**). This suggests that the dense phase pH environment is critical to the hydrolysis function of condensates.

### Programming IDP sequence to modulate the interfacial electric fields of condensates

To understand how the interfacial electric fields contribute to catalytic reactions, we explored whether we could design IDP sequences to modulate the surface charge density of condensates without changing dense phase environments. We hypothesized that the surface charge density of condensates will define the strength of the interfacial electric field ^59,60^, thereby modulating the catalytic behaviors. Similar to colloidal particles in a solution containing electrolytes ^61,62^, the first layer of ions within the Stern layer ^63^, which is governed by the surface charge of the colloid particle, will organize the assembly of the second layer of ions that screens the first layer within the shear plane. This gives rise to the electric double layer that defines the strength of the interfacial electric field.

We hypothesized that if certain regions of IDPs preferentially accumulate at the interface of the condensate, then we should be able to tune the surface charges of condensates ^64^. This would allow us to alter the interfacial ion organization (**Figure 2A**). To this end, we first employed a coarse-grained simulation approach using LaSSI^65^, which is a Monte-Carlo simulation engine that has been used to model phase behaviors of low-complexity sequences and the internal organization of condensates ^28,66^. The simulations helped us explore if there is a defined region of RLP_WT_ that has a higher preference of localizing to condensate surface. The repetitive nature of the RLP sequence mediates strong homotypic interactions. We focused our attention on the N-terminus (Ser-Lys-Gly-Pro), which lacks sticker residues ^67,68^ and is distinct from the repetitive main sequence. We hypothesized that the N-terminus should interact weakly with repetitive elements within the dense phase and that the charged Lys residue in the N-terminus might drive a preference for the N-terminus to localize to the condensate surface.

**Figure 2.**
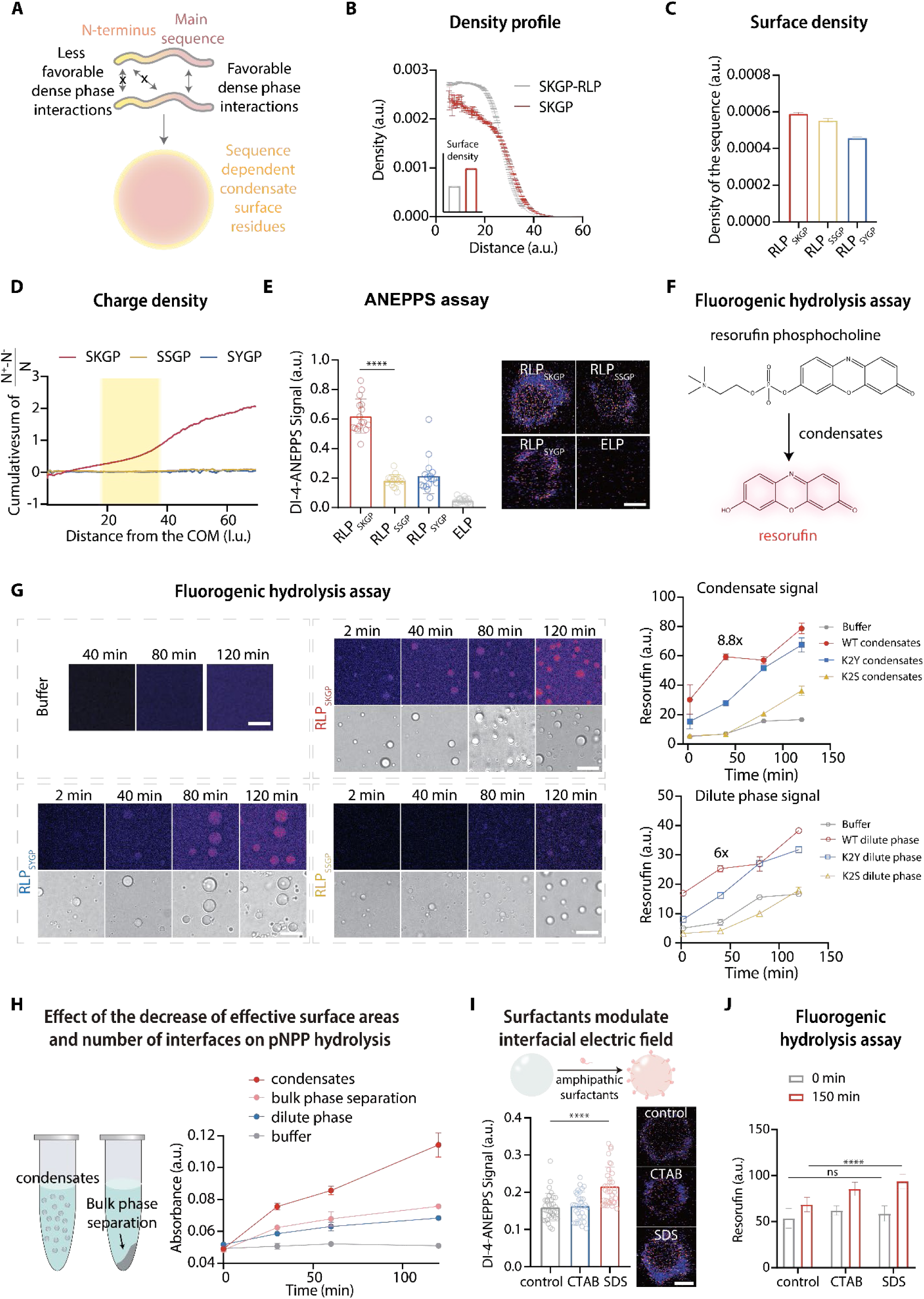
Decoupling the contribution of the dense phase chemical environment and the interfacial electric field to the catalytic function of condensates. **A,** Schematic showing that regions of an IDP can have different intrinsic preferences for localizing to the dense phase versus localizing to the interfaces of condensates. **B,** Radial density plots of the N-terminal region (first four residues) and full-length protein from simulations of the full-length RLP_WT_. Each data point represents mean ± SE. Inset shows the comparison of the density of the N-terminal sequence and the full-length sequence at the interface of the condensates based on the radial density plots. Bar graph shows the mean ± SE. **C,** Data from the radial density plots in panel B were used to compute the interfacial densities of the N-terminal motifs in different condensates. **D,** Comparison of the cumulative density of net charge computed along the radially resolved coordinate from the center of mass of each condensate for each of the systems featuring N-terminal sequences: Ser-Lys-Gly-Pro, Ser-Ser-Gly-Pro, and Ser-Tyr-Gly-Pro. Bar graph shows the mean ± SE. **E,** Evaluation of the interfacial electric field of different sequences (SKGP-RLP, SSGP-RLP, SYGP-RLP, ELP) using ratiometric DI-4-ANEPPS fluorescence assay. The insets show the sample ratiometric images of condensates (at similar size) with DI-4-ANEPPS. Scale bar, 1 µm. **F,** Schematic showing how fluorogenic hydrolysis reaction of resorufin phosphocholine generates fluorescent resorufin. **G,** Comparison of the resorufin signal of the fluorogenic assay using different condensates formed by RLPs with different N-terminal sequences namely, SKGP-RLP, SSGP-RLP, and SYGP-RLP. The excitation was set at 550 nm with a WLL laser, and the emission detector was set at 570-600 nm on a HYD detector. The highest fluorescence signal within a 15 μm z range was identified for quantification of the resorufin signal. The grey line represents the spontaneous hydrolysis of resorufin phosphocholine in the same buffer solution. Scale bar is 5 μm. Each data point represents mean ± SE. **H,** Schematic illustration of the experimental design for creating bulk phase separation by spinning down the condensates to reduce the effective surface areas and number of interfaces. Additionally, the pNPP hydrolysis assay analysis was conducted to evaluate the difference in hydrolytic capability between dense phases in the form of condensates and bulk phase separation. **I,** ANEPPS assay analysis of the change of interfacial electric field of RLP_SKGP_ condensates after being treated by charged surfactants (CTAB and SDS). The schematic illustration shows how amphipathic surfactants attach onto the surface of condensates. Scale bar, 1 µm. **J,** Fluorogenic hydrolysis assay based on resorufin phosphocholine analysis of the change of hydrolytic capability of RLP_SKGP_ condensates after being treated by charged surfactants (CTAB and SDS).

We performed simulations using the full-length protein and analyzed the resulting radial density profiles. This analysis focused on the localization preferences of the N-terminus versus the rest of the sequence. As expected, the N-terminus shows a decreased density in the dense phase (**Figure 2B**) and an increased density at the interface when compared to either the C-terminus or the full protein. This observation suggests that the specific sequence of the N-terminus (Ser-Lys-Gly-Pro) may promote its localization to the interface.

Next, we made variants containing different N-terminal sequences by replacing the charged residue, Lys, in the N-terminus sequence with a Ser or Tyr. This sets up a hierarchy whereby the wild-type (SKGP) N-terminus, which contains a Lys residue, would have the weakest interaction with the main repeat, followed by the N-terminus with a Ser residue (SSGP/K2S), and the N-terminus with a Tyr residue (SYGP/K2Y), which should have the strongest interaction with the main repeat. Indeed, we observed that the density of the N-terminal sequence motif in the interface was significantly diminished for the variants when compared to the WT sequence, with the N-terminus containing a Tyr showing the lowest preference for localization to the interface (**Figure 2C**). This finding confirms that the relative interaction strengths between distinct sequence regions can affect the spatial preferences of each region within a condensate ^66^. This also provides a fundamental ability to program the interfaces of condensates without disrupting interactions within dense or dilute phases, thereby allowing us to isolate contributions from the interfacial electric field to catalytic behaviors.

To understand how sequence variations contribute to charge asymmetry, which is the key to set up a potential gradient, the radial density profiles were computed for positively charged and negatively charged residues in the system to analyze the net charge across the radial coordinate from to the centers of mass of different condensates in the simulations. The cumulative sum of the net charge radial density profile compared to the radial distribution of all beads in the system shows whether there is an accumulation of positively charged or negatively charged beads at specific regions, within the condensate, in the interface region (shaded), and in the dilute phase. (**Figure 2D**). Indeed, for the WT sequence, the concavity of the profile in the interface shows accumulation of positively charged residues. For K2S and K2Y, we did not observe any significant changes in the accumulation of net charge across the radial distances from the center of mass of the droplet. These simulation results suggest that WT sequence generates the highest interfacial potential gradient.

We tested the predictions from simulations using experiments that investigated the effects of distinct sequences on the interfacial electric field and the catalytic behaviors of the resulting condensates. We hypothesized that removal of the charged residue at the interface of condensates modifies surface electrostatics and ion alignments ^69,70^, thereby changing the strength of the interfacial electric field and catalytic capabilities. To ensure that the modification of a single-amino acid residue at the N-terminus does not affect the internal properties of condensates, we first measured the dense phase concentrations of these condensates and did not observe significant differences (**Figure S3B**). This suggests that the homotypic interactions between the main repetitive sequences stabilize the dense phase. Next, we probed the pH of these condensates using a previously established method ^14,27^ and found that the N-terminus sequence does not significantly affect the dense phase pH (**Figure S3C**), confirming that the main repetitive sequences determine the chemical environments of the dense phase.

To investigate how the interfacial properties of condensates change due to single mutations in the N-terminus, we used a ratiometric DI-4-ANEPPS dye, which undergoes a structural change and fluoresces upon experiencing a local electric field ^14,71,72^. The ratiometric feature of ANEPPS allows us to quantify the electric field strength without considering the concentration-dependent effect of the dye ^71^. We compared condensates of similar sizes and found a noticeable difference in the interfacial signal between the WT, K2S, and K2Y sequences. The interfacial potentials are strongest for WT, followed by K2Y, and weakest for K2S, with an over 5-fold lower signal generated by K2S (**Figure 2E**). These observations align with the simulation results and suggest that the interfacial alignment of ions, which can be modulated by surface electrostatics dictated by the protein sequence, will perturb local ion density and modulate the strength of the interfacial electric field.

### Interface-dependent catalytic activity of condensates

Sequences that encode the same dense phase chemical environments of condensates while enabling distinct interfacial features enable the quantification of contributions of interfacial electric fields to the catalytic functions of condensates. To study this feature at a single condensate level, we designed a fluorogenic assay using resorufin phosphocholine as the substrate. Hydrolysis of this substrate removes the phosphocholine and yields a fluorescent resorufin (**Figure 2F**). We tracked the catalytic behaviors of condensates formed by different sequences as a function time (**Figure 2G**). Specifically, we compared the resorufin signal in different condensates of similar size (r =1-3 μm). The resorufin signal of condensates formed by RLP_WT_ was significantly higher than that of condensates formed by the mutants K2S and K2Y. The most significant difference was recognized at 40 min with an 8.8-fold catalytic product difference between the condensates of RLP_WT_ and the condensates formed by K2S (**Figure 2G**). This signal difference between different types of condensates possessing similar dense phase environments suggests that the resorufin is not specifically partitioned into the condensates. Instead, the trends in resorufin signals, which reflect the catalytic activity of different condensates, correlate with their differences in interfacial potentials.

Next, we assessed whether the dilute phase environment is critical to the catalytic behavior of condensates. To this end, we modulated the bulk phase environmental pH of solutions containing RLP condensates. This experiment was motivated by previous discoveries showing that the dense phase pH of RLP_WT_ condensates does not shift significantly upon a change in the bulk phase pH ^14^, which suggests that the dense phase interactions define the interior microenvironments. With the fluorogenic assay, we found that compared to condensates prepared in solutions that are more basic, a considerably higher fluorescent signal was generated from RLP_WT_ condensates in solutions that are more acidic (**Figure S3D-F**), even though a basic bulk environment is favorable for the hydrolysis of resorufin phosphocholine. We reasoned that decreasing the bulk phase pH of the solution containing RLP_WT_ condensates (with an alkaline dense phase) would generate an increased interphase pH gradient between the dilute and dense phases ^14^. This, in turn, should lead to a stronger interfacial electric field driven by the larger potential gradient. Taken together with data for ELP versus RLP condensate and RLP condensates with distinct sequences or under distinct environments, our observations suggest that the interphase electrochemical properties defined by differences between coexisting dilute and dense phases, and the interfacial electric fields are both critical to the catalytic behaviors of condensates (**Figure S3G**).

To investigate the specific contributions of the interfaces of condensates, we designed two different assays to isolate the contribution of interfaces of individual condensates on catalytic functions. First, we used bulk phase separation to generate a single interface that separates the dilute and the dense phases (**Figure 2H**). We implemented the pNPP assay to study the contributions of individual phases to the hydrolysis reaction in bulk solutions. Compared to the condensate solution, in which many interfaces exist, the solution that was separated into bulk phases showed 2.44-fold decreases of the catalytic behavior (**Figure S3H**). By extracting the dilute phase, we found a slightly higher level of the catalytic activities comparing to that of the buffer solution, which might be attributed to the existence of clusters in subsaturated solutions ^73–75^ (see Discussion). Second, to isolate the interfacial effect, we implemented anionic and cationic amphipathic surfactants, sodium dodecylsulfate (SDS) and cetyltrimethylammonium bromide (CTAB), to perturb the surface charge conditions of the condensates (**Figure 2I)**. This should affect the alignment and distributions of surface ions ^76^, thereby affecting the interfacial field of the condensates. We verified the change of interfacial field of individual condensates upon the addition of surfactants with the ANEPPS assay (**Figure 2I**). We then implemented the fluorogenic hydrolysis assay and compared the change of generated resorufin signal of these condensates with similar size during the same amount of the reaction time. We found that compared to the condensates with smaller electric fields, the condensates with added SDS, which showed a substantially larger interfacial field, significantly enhanced the hydrolysis reaction (**Figure 2J**). These results collectively confirm that the interface of condensates is crucial to their catalytic performance.

### Condensates regulate catalytic behavior by creating heterogenic water environments

We next set out to study condensate-dependent water activity, which should directly contribute to the hydrolysis reactions. The abundance of free water molecules that can form hydrogen bonds with solutes or substrates has also been correlated with the rates of catalytic reactions ^77,78^. In addition to participating in the hydrolysis reaction, water molecules also contribute to the establishment of interfacial electric field. In addition to the surface electrostatics that define the interfacial electric fields as described by the Gouy-Chapman-Stern formalism ^79^, the total field generated by the two-phase system can also be influenced by the solvent dipoles that are mediated by solvent-solute interactions ^80^. An example of this is the Onsager reaction field at an electrochemical interface ^81^. We reasoned that as the polarity of water molecules and the interfacial electrostatic field are always coupled together, one might affect the other or *vice versa*. Therefore, different water activities might exist in different types of condensates, which could be the key factors contributing to the hydrolysis reactions.

To test for the possibility of spatially varying water activities, we employed stimulated Raman scattering (SRS) microscopy ^82–87^ to study the hydrogen-bonding environment by imaging the local water status in a label-free manner (**Figure 3A**). Harnessing the hyperspectral chemical imaging capability and submicron resolution of SRS microscopy, we mapped the authentic water vibrational signatures within the O-H stretching window for the three distinct RLP condensates (**Figure 3A)**. The Raman signature of water O-H stretching is particularly sensitive to the hydrogen-bonding environment, in which three distinct types of water molecules can be distinguished through their frequency regions (**Figure 3A**) ^88–90^. In general, the red-shift of the OH frequency corresponds to a stronger hydrogen bonding network with more hydrogen bonds between water molecules, while the blue-shift means more defects in the hydrogen-bonding network with fewer hydrogen bonds between the water molecules. By quantifying the intensity ratio obtained from the spectrum of different types of hydrogen bonding, for all the three sequences (WT, K2S, K2Y), we observed spatial heterogeneity of the local water environment in the condensate systems, where higher populations of free water and lower populations of tetrahedral water were observed at condensate surface (**Figure 3B,3C** and **Figure S4**). We further quantified the overall strength of the hydrogen-bonding network by calculating the average O-H frequency 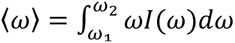, where the corresponding intensity is normalized to the unit area between *ω*_1_=3100 cm^−1^ and *ω*_2_=3800 cm^−1^. The maps of the averaged frequency 〈*ω*〉 show similar patterns of spatial heterogeneity of the local water environment, where 〈*ω*〉 at the condensate interface shows a distinct blue-shift, indicating a weakened hydrogen bonding network at condensate interface (**Figure 3D**). This is reflected as a pattern of anisotropy, which is likely due to the alignment of the water dipole at the condensate interface modulated by the surface electrostatics and hydrogen bonding interactions.

**Figure 3.**
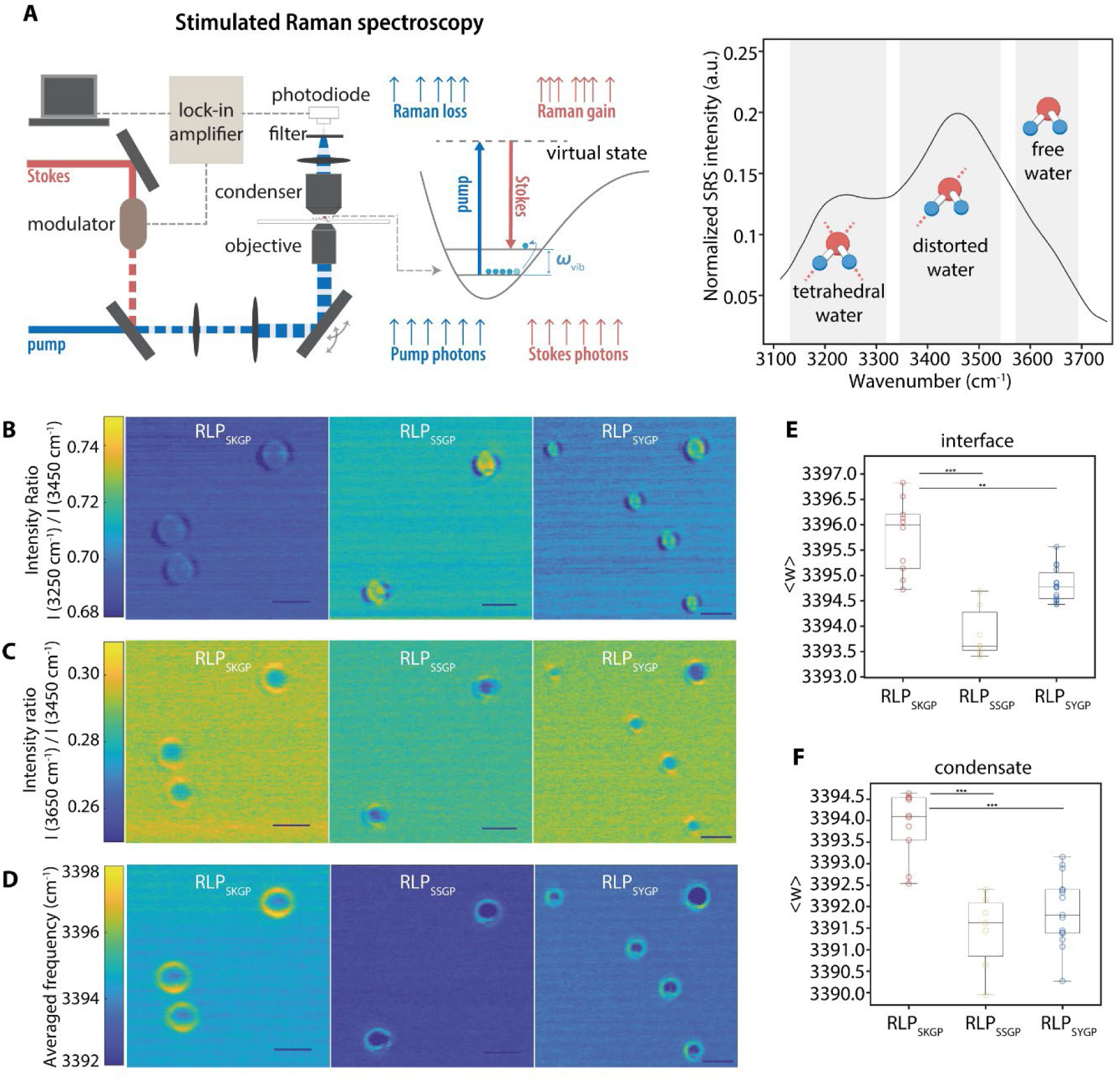
Characterization of water environments using hyperspectral stimulated Raman Spectroscopy. **A,** SRS microscopy illustration. The pump laser beam and modulated Stokes laser beam are combined and focused onto the sample. When the energy difference between the pump and the Stokes laser matches the vibrational energy (ω_vib_) of the chemical bond of interest, the vibrational transition is accompanied by stimulated Raman loss (SRL) on the pump beam and stimulated Raman gain (SRG) on the Stokes beam. SRL on the pump beam is detected by the photodiode and extracted by the lock-in amplifier for SRS imaging. **B,** Hyperspectral SRS mapping of water heterogeneity in condensates with ratiometric SRS images (I(3250 cm^−1^) /I(3450 cm^−1^) shows the relative abundance of tetrahedral water to distorted water. Scale bar, 3 µm. **C,** Hyperspectral SRS mapping of water heterogeneity in condensates with ratiometric SRS images (I(3650 cm^−1^) /I(3450 cm^−1^) shows the relative abundance of free water to distorted water. Scale bar, 3 µm. **D,** Hyperspectral SRS mapping of water heterogeneity in condensates with images shows the averaged OH frequency <ω>. Scale bar, 3 µm. **E,** Quantitative analysis of averaged OH frequency <ω> for the interfaces of different condensates. **F,** Quantitative analysis of averaged OH frequency <ω> for the dense phase of different condensates. Unpaired T-test shows the p-value.

To explore whether the sequence-dependent catalytic behavior is reflected in the water activity, we compared the blue-shifted 〈*ω*〉 values at condensate interfaces of three distinct condensates by comparing condensates of similar sizes. We observed that the averaged frequency 〈*ω*〉 at WT condensate interface is significantly higher than those of K2S and K2Y condensates (**Figure 3E, 3F**), suggesting a higher abundance of free water molecules. This aligns with previous work suggesting that weakened hydrogen bonding networks or increased free water/bound water ratios facilitate the participation of water molecules in the catalytic reactions ^77,78,91^. This observation further supports the view that the interfacial properties of condensates can be modulated by a single mutation in the N-terminus. Compared to K2S and K2Y, the significantly blue-shifted 〈*ω*〉 values of the WT condensate both inside the condensate and at the condensate interface suggest that the charged amino acid induced considerable disruption to the hydrogen bonding network of the water microenvironment. The observed chaotropic properties of the lysine residue are consistent with the reported effect from its interaction with water molecules in competition with the water hydrogen bonding network ^92^. These results collectively confirm that the interfacial electrostatic properties can affect the catalytic performance, which involves a synergistic contribution from local water activity modulated by charge-dependent hydrogen bonding and interfacial electric fields.

### Condensate-dependent hydrolysis is generalizable

We next asked if condensates formed by naturally occurring IDPs also possess catalytic functions. To explore whether the catalysis of hydrolysis by condensates might be a generic feature, especially if the IDPs feature ionizable residues, which are essential for mediating a great extent of asymmetric ion distribution in the co-existing phases ^22^, we reconstituted different condensates formed by IDRs of the Dead-Box helicase 4 (Ddx4) from nuage bodies ^93^, the prion-like low complexity domain of hnRNP A1 (A1-LCD) from stress granules ^68^, and the nucleolar protein Ly-1 antibody reactive (LYAR) ^11^. We used the pNPP assay to evaluate whether condensates formed by naturally occurring IDPs can catalyze reactions. We found that compared to the buffer solution and the subsaturated protein solution, solutions containing condensates formed by Ddx4 and LYAR were able to catalyze the decomposition of pNPP, while A1-LCD condensates showed minimal functionality as catalysts (**Figure S5A**). Ddx4 condensates showed the most significant activation with an over 4-fold increase of catalytic products after 150 min incubation at room temperature. The distinct catalytic behaviors of condensates formed by Ddx4 and LYAR versus A1-LCD can be explained by the high fraction of charged residues and polyampholytic character of the IDRs of Ddx4 ^93^ and LYAR ^11^. In contrast, the A1-LCD has a very low fraction of charged residues ^68^.

We also used the pNPP assay to explore whether facsimiles of native condensates, which are complex coacervates formed via heterotypic protein-RNA interactions could mediate catalytic functions. For this, we selected condensates formed by nucleophosmin 1 (NPM1) and mature rRNA ^11^ (mat-rRNA), which are also defined by an interphase pH gradient ^11^. Compared to the solutions containing only NPM1 protein, mat-rRNA, and buffer, the solution containing the NPM1-rRNA condensates was able to accelerate the decomposition of pNPP (**Figure S5A**). These observations suggest that the inherent catalytic functions of condensates are encoded in the molecular grammars of the biomolecules and the partitioning features defined by the driving forces for condensation. Our findings regarding the catalytic activities of NPM1 + mat-rRNA condensates are likely to have broad implications for activities at the interface between granular components of nucleoli and the nucleoplasm.

### Hydrolysis reactions for cellularly relevant substrates

We next asked whether condensates can modulate different types of hydrolysis reactions relevant to cellular processes. Specifically, we investigated the hydrolysis of disaccharides, peptides, NTPs, dNTPs and nucleic acids with RLP_WT_ condensates using mass spectroscopy and fluorogenic assays (**Figure 4A**). These substrates are common chemicals undergoing hydrolysis reactions critical to metabolic cycles ^94^. For disaccharides and peptides, we studied the hydrolysis of lactose and N-Succinyl-Ala-Ala-Ala-*p*-nitroanilide. We discovered that condensates hydrolyzed lactose into glucose and galactose while we did not find any product in the solution without condensates (*c*_RLP_<*c*_sat_). For succinyl-Ala-Ala-Ala-p-nitroanilide hydrolysis, compared to the solution without condensates, we found that condensates improve the hydrolysis rate over 15.8-fold (**Figure 4B, 4C** and **Figure S5B, S5C**). For the hydrolysis of NTP and dNTP, we found that condensates are capable of hydrolyzing all types of NTPs, generating dephosphorylated products with relative abundances exceeding 50%, whereas NTPs in solution without condensates remain intact in the same reaction condition. Similar results were observed for dNTPs (**Figure 4D** and **Figure S5D, S5E**). Lastly, we studied the hydrolysis of nucleic acids by employing a single-stranded DNA (ssDNA) containing a quencher pair, 6-FAM and BHQ-1, at its 5’-and 3’- ends (**Figure 4E**), the hydrolysis of which breaks the phosphodiester bond and leads to the release of fluorescent 6-FAM. We found time-dependent increase of fluorescence only in the solution containing condensates with a stable background signal. The capability to hydrolyze nucleic acid by condensates was further verified with a supercoiled plasmid (**Figure S5F**). These observations suggest that the capability of hydrolysis of condensates is generalizable to different biologically relevant substrates.

**Figure 4.**
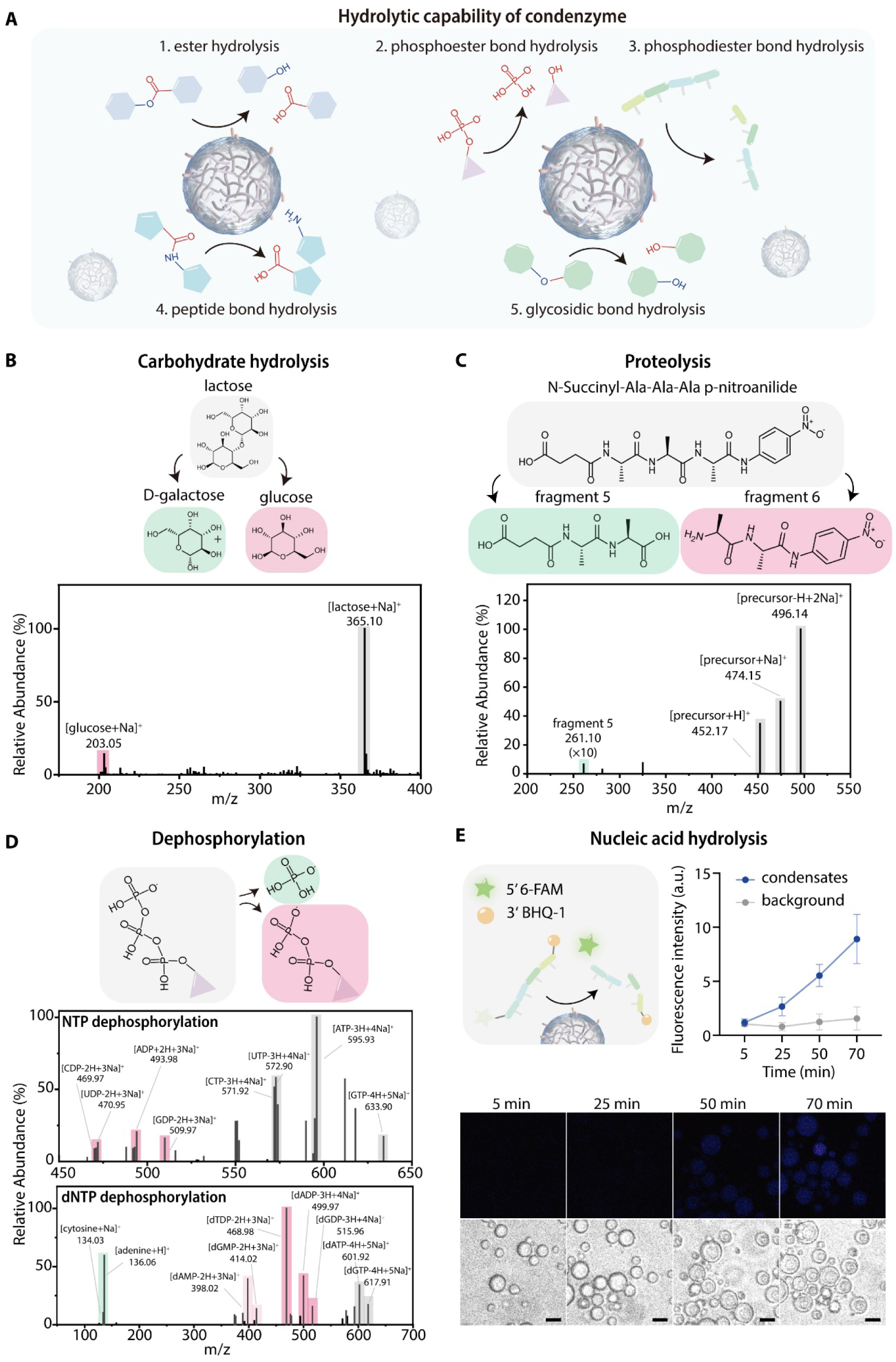
Characterization of the generality of hydrolytic capability of condensates towards disaccharide, peptide, NTPs, and nucleic acid. **A,** Schematic demonstration of the hydrolysis reactions catalyzed by the RLP condensates. These hydrolysis reactions include esterolysis, dephosphorylation, nucleic acid hydrolysis, peptide hydrolysis, and disaccharide hydrolysis. **B,** Mass spectroscopy assay analysis of the hydrolysis of glucose by the condenzyme, which results in the generation of D-galactose and glucose. **C,** Mass spectroscopy assay analysis of the hydrolysis of peptide by the condenzyme. The results indicate the cleavage of peptide bond by the condenzyme, resulting in the formation of peptides with smaller molecular weight. **D,** Mass spectroscopy analysis assay of the hydrolysis of NTP and dNTP by the condenzyme. The results indicate that condenzyme can hydrolyze all NTP and dNTP into phosphate and different dephosphorylated products. . **E,** Mass spectroscopy assay analysis of the hydrolysis of ssDNA by the condenzyme. The increase of fluorescent intensity by the hydrolyze fluorogenic probe (6-FAM-ssDNA-BHQ-1) indicate the hydrolytic capability towards ssDNA. Scale bar, 5 µm.

### Condensates can fully decompose ATP

Considering the significance of hydrolysis of ATP in energy generation and cellular signaling, we next focused on studying the hydrolysis of ATP by condensates (**Figure 5A**). ATP hydrolysis is a process typically mediated by ATPases ^95^, in which ATP could be decomposed to ADP and AMP through dephosphorylation reactions ^96^. We first quantified the hydrolysis of ATP using a bioluminescence-based ATP quantification assay (see **Figure S6A** for calibration). After incubating condensates with ATP for 30 minutes at room temperature, we found that over 95% of the ATP was decomposed by condensates (**Figure S6B**). To analyze the products of the decomposition reaction of ATP, we employed high resolution linear ion mass spectroscopy. Surprisingly, we not only found the existence of ADP and AMP, but we also detected distinct peaks at *m/z* ratios of 134.05, 105.02, 75.00 that correspond to adenine and different carbohydrates (**Figure 5B**). This surprising finding suggests that the RLP_WT_ condensates not only facilitate the dephosphorylation of ATP; they also contribute to the bond breakage of adenosine to form adenine and different carbohydrates. This bond breakage might be attributed to the free radicals generated by condensates through their spontaneous redox activities ^14,97^. This observation points to an important role of condensate-based catalysts in generating cellular energy and it points to the critical importance of interfacial electric fields ^14,56^, which can provide the energy source for such reactions. These results further highlight the fact that condensate-specific microenvironments and interfacial properties jointly contribute to the ability of condensates to catalyze reactions.

**Figure 5.**
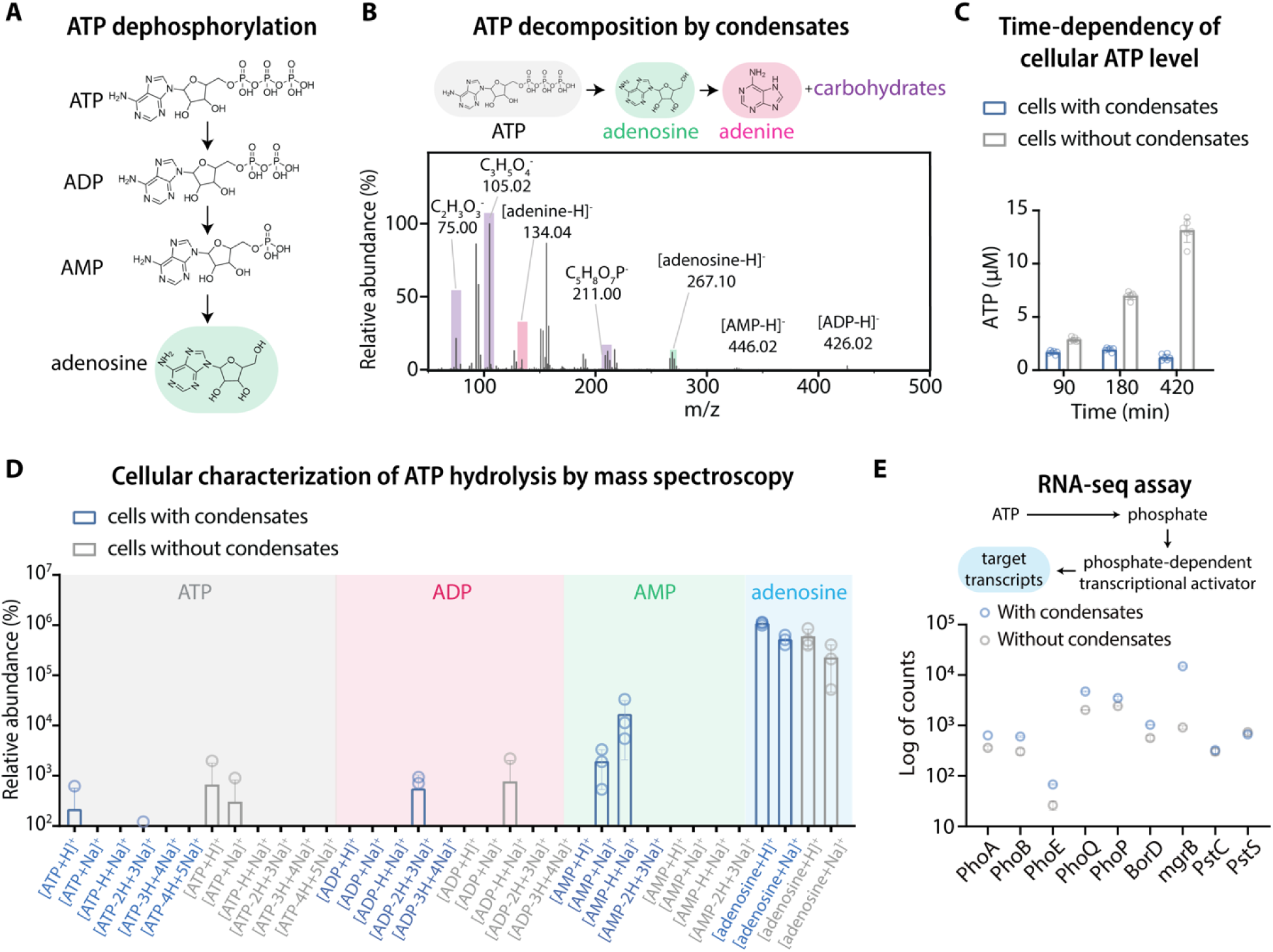
Characterization of the hydrolytic capabilities of the condenzyme in cells and their biological implications. **A,** Schematic illustration of the dephosphorylation of ATP into ADP, AMP, and adenosine. **B,** Mass spectroscopy assay analysis of the in vitro hydrolysis of ATP by the RLP condensates into ADP, AMP, adenosine, adenine, and carbohydrates. **C,** Cellular ATP quantification assay analysis of the change of ATP level in cells with and without condensates at different time point. **D,** Mass spectroscopy assay analysis of the cellular ATP, ADP, AMP, and adenosine level in cells with or without condensates. **E,** RNA-seq assay analysis of the difference in ATP dephosphorylation-related gene expression in cells with or without RLP condensates. N=2 independent experiments.

Next, we asked if synthetic condensates can function as catalysts in living cells. We transformed *E. coli* BL21 cells (DE3) with an isopropyl β-D-1-thiogalactopyranoside (IPTG) inducible pET24 plasmid encoding the RLP_WT_ under a T7 promoter ^43^. We implemented a cellular ATP assay to probe the change in ATP concentration in cells with or without condensates during their exponential growth phases. The experimental condition to create cell with or without condensates has been well-calibrated in a previous work ^98^. To decouple the effects of inducing protein expression, we compared ATP levels in cells with or without induction of the expression of a cyan fluorescent protein based on the same plasmid backbone. We did not detect a decrease in ATP level upon protein expression (**Figure S6C**). This aligns with a previous observation that the induction of pET based vectors in BL21(DE3) results in the accumulation of ATP and metabolites caused by excessive energy generation from the expression system ^99^. To understand the effects of condensates on cellular ATP, we cultured cells with or without condensates in the rich 2 x YT medium and tracked cellular ATP levels. After three hours of culturing, the cells containing condensates showed a 3.6-fold lower ATP level than those without condensates (**Figure 5C**), which was further enhanced after 7 hrs of culturing.

Next, we leveraged the leaky expression nature of the T7 promoter to track the correlation between condensate formation and the decrease of cellular ATP levels, thereby fully excluding the possible metabolic burden caused by overexpression on the consumption of ATP ^100,101^. We applied differential interference contrast imaging to track condensate formation and correlated the results with the time-dependency of ATP levels of cells grown in the nutrient-rich 2 x YT medium. We observed a gradual increase of ATP levels alongside exponential cellular growth before condensate formation and a sharp decrease (>30%) of ATP levels at a time point that aligns with condensate formation (**Figure S6D and S6E**). This process was studied within a short period and in a rich medium, thereby eliminating the possibility of nutrients limitations.

Finally, we implemented non-targeted mass spectroscopy to quantify the products of ATP hydrolysis in cells with or without condensates at their same exponential growth stage. We found that the products underlying the ATP hydrolysis pathway were substantially upregulated in cells with condensates compared to cells without condensates (**Figure 5D**). These observations confirm that the catalytic functions of condensates remain operative in living cells and point to a role for condensates as modulators of cellular energy sources.

### Condensate-mediated ATP decomposition activates phosphate-dependent cellular pathways

To evaluate whether the condensate-mediated ATP decomposition can activate specific cellular pathways, we implemented RNA-sequencing to quantify the targeted transcriptome correlated with the consequences of ATP decomposition. We specifically assessed free phosphate-dependent cellular pathways in *E. coli* ^102–106^, in which free phosphate binds to the transcription factor and activates downstream gene expression. We prepared cells with or without condensates from log-phase culture and analyzed them using RNA-sequencing. We found that the gene clusters driven by phosphorylated PhoB and PhoP transcription factors ^104,107,108^ are significantly upregulated (p-value < 0.048) when comparing cells with condensates and cells without condensates (**Figure 5E**). Similarly, based on the same logic, we found that the fatty acid hydrolysis-dependent pathway was also upregulated in the cells with condensates (**Figure S6F**). We also compared the capability of ELP condensates on driving these pathways and found that though the capability of ELP condensates is not as noticeable as the RLP condensates, ELP condensates were still able to activate certain pathways (**Figure S6G**). Considering the complexity of intracellular environments, the phase transition of ELP in living cell can potentially mediate associate transition of other biomolecules, thereby setting up different conditions as tested in simple in vitro setting (see Discussion for detailed mechanism). This suggests that the catalytic functions of condensates generate global changes of cellular chemical landscapes that can potentially affect cellular pathways.

### Catalytic functions of condensates modulate intercellular signaling

The robust capability of condensates to modulate intracellular catalytic reactions suggests a potential role of condensates on cellular signaling through their inherent catalytic functions. To test this possibility, we first implemented two orthogonal sets of intracellular fluorogenic hydrolysis assays using fluorescein-di-beta-D-galactopyranoside (FDG) and fluorescein-diphosphate (FDP), the hydrolysis of which results in the generation fluorescein. The capability of RLP condensates to hydrolyze FDG and FDP probes was first verified *in vitro* (**Figure S7A-S7D**). Subsequently, we incubated the probes with cells with or without RLP condensates in their exponential phases at the same cellular density. After 30 min of incubation, compared to cells without condensates, we found over 100-fold increase of the fluorescein signal in the cells with condensates (**Figure 6B**). We also compared the catalytic behaviors of ELP condensates and RLP condensates in cells and found that ELP condensates showed limited catalytic activity (**Figure S7E&F**). These behaviors suggest condensates can transform heterogeneous signaling molecules in living cells, which can serve as a potential strategy to modulate inter-cellular signaling for synthetic biology ^109–113^.

**Figure 6.**
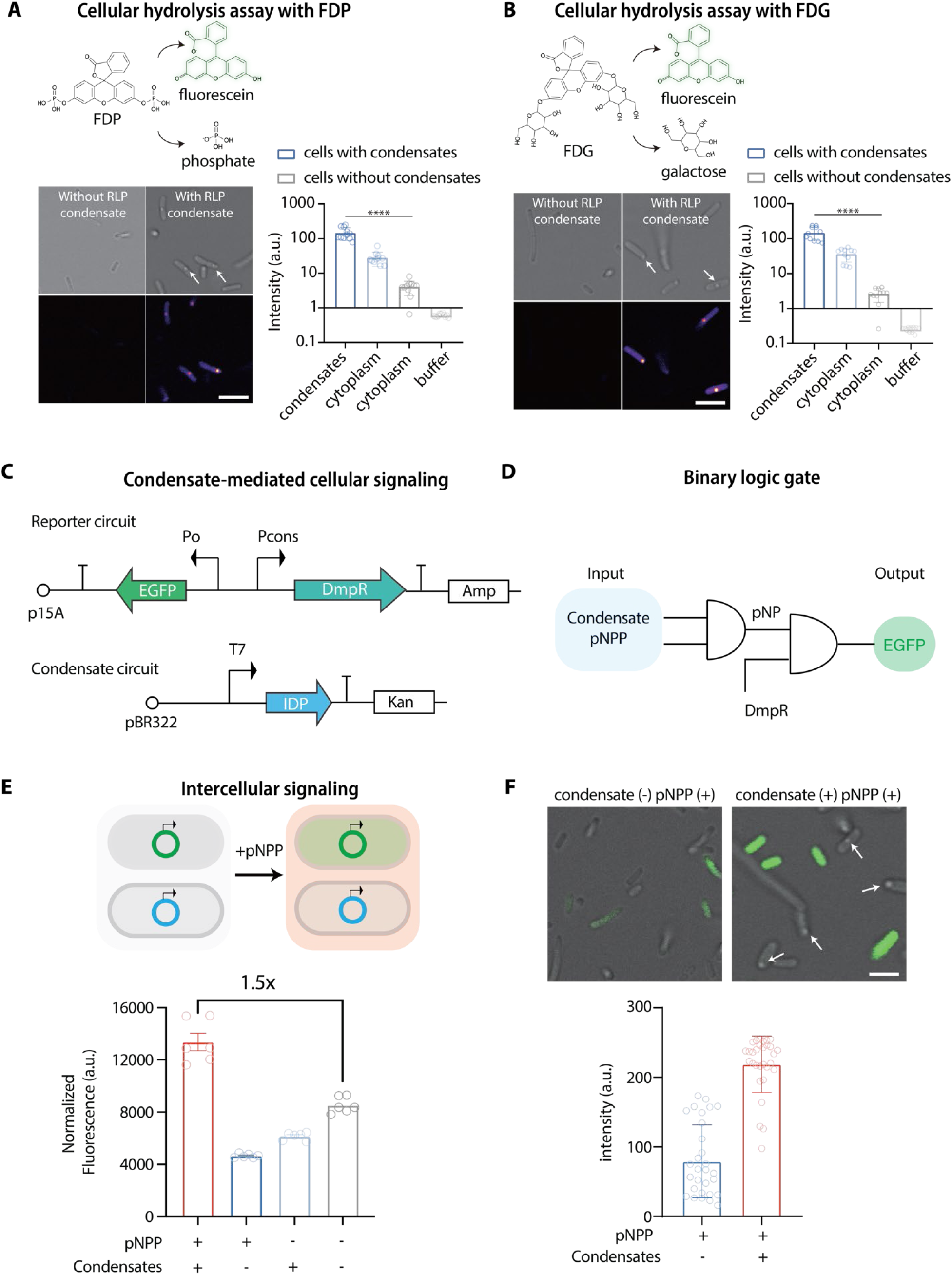
Catalytic functions of condensates mediate the activation of synthetic gene circuits in *E. coli*. **A,** FDP probe-based fluorogenic hydrolysis assay analysis of the hydrolysis capability of cells expressing RLP protein but not yet formed condensates and cells with RLP condensates. White arrows point to the condensates in the phase contrast imaging. Scale bar, 2 µm. **B,** FDG probe-based fluorogenic hydrolysis assay analysis of the hydrolysis capability of cells expressing RLP protein but not yet formed condensates and cells with RLP condensates. White arrows point to the condensates in the phase contrast imaging. Scale bar, 2 µm **C,** A reporter circuit was built using DmpR, a phenol activated transcription activator as a sensor, to drive the expression of EGFP, which serves a reporter protein for the cellular signaling. A condensate circuit was built to generate the cellular signaling event, which is mediated by condensates-mediated pNPP degradation to produce phenol. The coupling of these two circuits can be used to detect whether condensates have the capability to drive cellular signaling through degradation of pNPP from the culture medium. **D,** The designed gene circuit follows a binary logic gate cascade with the formation of condensates and the existence of pNPP as the inputs to generate pNP, which activates DMPR to drive the expression of EGFP protein as the output. **E,** Evaluation of the capability of condensates to mediate intercellular signaling with the condensate circuit transformed into the *E. coli.* BL21 (DE3) cells and the reporter circuit transformed into the *E. coli.* DH5α cells. The fluorescence signal was normalized to the cellular density. N = 6 independent experiment. Bar graph shows the mean ± SD. **F,** Confocal microscopy analysis of GFP expression in E. coli DH5α cells containing the reporter circuit, cocultured with pNPP and E. coli BL21 (DE3) cells with or without RLP condensates. White arrows point to the condensates indicated by the phase contrast imaging. Scale bar, 2 µm.

We next designed and engineered a synthetic gene circuit that can be activated by the catalytic product of condensate-mediated hydrolysis reactions. We elected to use the phenol product generated from the pNPP reaction to activate a gene circuit (**Figure 6C**). To connect the phenol generation with a cellular response, we implemented a σ_54_-dependent phenol-responsive transcription regulator DmpR ^114^, which specifically drives the activation of the phenol degradation pathway in bacteria ^115^, to regulate the expression of an enhanced green fluorescent protein (EGFP) (**Figure 6D**). This serves as the downstream reporter, thereby establishing a logical AND gate signaling cascade initiated by the chemical activity of condensates. We confirmed the activity of the DmpR circuit by monitoring the EGFP response in cell cultures with or without the phenol product. The cell culture with 100 μM phenol showed a 3-fold higher normalized EGFP signal compared to the culture without phenol (**Figure S7G**).

We explored whether the catalytic activity of condensates could be used to control different populations of cells based on intercellular signaling. We transformed the *E. coli.* BL21 (DE3) cells with the plasmid encoding the RLP_WT_ as the signaling cell and the *E. coli.* DH5α with the reporter circuit as the sensor cell. We then co-cultured the two populations of cells in one culture (**Figure 6E**). Specifically, we incubated log-phase BL21 (DE3) cells with or without condensates with DH5α cells in M9 medium containing pNPP in an equal volume fraction and tracked the activated EGFP signal. We found that only the BL21 (DE3) cells with condensates could activate the EGFP signal in the DH5α cells (**Figure 6F**). To test the robustness of the signal, we varied the volume fraction between the BL21 (DE3) cells with condensates and the DH5α cells under a fixed total cell density. We found that increasing the fraction of BL21 (DE3) cells in the multicellular population could further increase the EGFP signal (**Figure S7H**), even though the absolute quantity of DH5α cells was decreased. This observation suggests that the rate limiting step for the intercellular signaling was the activation of gene circuits. Our results show that condensates can mediate intercellular signaling, thus establishing that the catalytic potentials of condensates can directly affect cellular population. This provides a new design principle for the construction of synthetic biological networks using the sustained chemical functions of biomolecular condensates. It also highlights the fact that condensates are unlikely to be just passive storage depots that are in any way incidental to cellular functions ^116^. Instead, we reason that catalytically active nature of condensates have a direct influence on the regulation of biochemical landscapes in cells.

## DISCUSSION

In this work, we showed that biomolecular condensates can catalyze hydrolysis reactions. Our data are consistent with sequence-dependent dense phase chemical environments and condensate-specific interfacial electric fields being the defining features of condensates ^11,12,14,15,22,27,117^ that enable them to function as mesoscale functional enzymes. Accordingly, we refer to condensates that lack biomolecules with intrinsic catalytic activities but show abilities to function as catalysts to be “Condenzymes”. In enzymes, the creation of unique catalytic environments is made possible by the chemistry that is present in active sites, ^40,56^; for Condenzymes, the electrochemical environments and active interfacial electric fields at the interfaces of condensates, which are encoded in the solvent environments defined by the condensed biomolecules through phase transition, drive their catalytic behaviors.

A diverse set of cellularly relevant molecules can be hydrolyzed by condensates, including disaccharides, peptides, NTPs, and nucleic acids. Each of these substrates involve different types of chemical bonds available for nucleophilic attack by water molecules. A surprising finding is that the catalytic function of condensates goes beyond hydrolysis. The RLP-based condensates can catalyze the full decomposition of ATP. In sharp contrast to the hydrolysis of ATP by ATPase, which catalyzes the decomposition of ATP to ADP or AMP, the condensates we studied are capable of decomposing ATP all the way to adenine, which requires the removal of the ribose sugar molecule from adenosine. This appears to be realized by free hydroxyl radicals generated by the ability of condensates to drive spontaneous redox reactions ^14^. The free radicals can lead to the hydrolytic release of ribose ^118^. A recent study also showed that the redox environment of condensates can regulate protein activity in condensates by mediating disulfide bond formation ^119^. These findings emphasize the critical importance of understanding the role of free radicals on the chemical functions of condensates and also the role of free solvated electrons generated by redox reactions ^14^.

Using a synthetic biology approach^120^ as a proof-of-principle, we demonstrated that the catalytic functions of condensates are prevalent in living cells. Moreover, such chemical functions are robust enough to lead to global effects in a cell and in a population of cells. These findings further highlight the fact that condensates are not chemically inert storage depots or as entities that are incidental to cellular functions ^116^. Instead, they are biochemically active. We deployed a series of IDPs of native proteins including those from Ddx4, hnRNP A1 and LYAR. These IDPs are known as the drivers of phase separation for the formation of different condensates in mammalian cells. We found that the condensates formed by highly charged IDPs including the RLPs, and IDRs from Ddx4 and LYAR were robust in catalyzing reactions, while condensates formed by IDPs with lower charge contents, such as the A1-LCD and the non-charged ELP were less capable of catalyzing reactions. This can be rationalized by the possibility that charge contents of IDPs correlates with the extent of asymmetric ion distribution upon phase transition ^121–129^. This conjecture is supported by the active catalytic behavior of the condensates formed by complex coacervation between NPM1 and mat-rRNA.

Asymmetrical partitioning of solution ions across the coexisting phases defines a chemical environment for the dense phase that is distinct from the dilute phase ^11,12,22^. This gives rise to interphase electric potentials that are essential for generating interfacial electric fields ^11,12,22^. Given that native condensates, especially RNP granules are enriched in charged biomacromolecules ^3,4,37,38,130^, such as RNAs and oppositely charged proteins, the asymmetric partitioning of ions and protons will set up distinct microenvironments and interfacial electric fields that provide the impetus for the biochemical activities of condensates ^14,131,132^.

Given the ubiquitous nature of condensates in living cells, we propose that such functions of condensates can be critical to understand biogenesis and chemical homeostasis. Biogenesis requires cascading biochemical reactions thus requiring a chemical sink. In this case, the spontaneous catalytic activity of condensates can initiate the cascade by defining a continuous baseline chemical activity. Further, hydrolysis reactions can diversify the chemical landscapes and provide thermal energy to drive other reactions, which are also favorable for the increased chemical complexity during biogenesis. Chemical homeostasis, which suggests the maintenance of stable levels of chemicals in living cells, requires balanced consumption and production of chemicals ^133^. Such process can be achieved through chemical oscillation ^134^, which requires a constant negative feedback to maintain the chemical driving force. The capability to break chemicals through hydrolysis or radical-dependent reactions of condensates can ideally enable negative feedback to maintain a stable chemical synthesis and consumption rate. Such driving forces in the case of transcription and translation condensates might serve as the key factor maintaining stability and control over gene regulatory circuits.

### What are the factors encoding the catalytic functions of condensates?

A key feature driving the catalytic reaction is the interfacial electric field, which is the electric double layer defined as the Gibbs dividing plane in the interphase region where charge separation occurs ^23,135–137^. First, under electrochemical potential equilibrium, the differential partitioning of ions sets up an electric potential gradient between phases thus enabling the charge asymmetry required for setting up the double layer ^14,22,26,27,98,123,137,138^. In the case of biomacromolecule phase separation, the ion gradient can be established through ionizable residues on the biomacromolecules; however, the hydration preferences of the backbones of biomacromolecules can alter local water environment, which in turn can affect the partitioning of ions based on the free energy of hydration of ions ^139^. Therefore, in theory, any kind of asymmetry caused by phase transitions^23^, such as differences in water structure, water abundance or ion abundance between phases or even limited chain flexibility, should be able to create a solvent environment that can set up an electric potential gradient ^98^. Second, the presence of a physical surface in the solution results in anisotropic forces acting on the solvent ^61,140,141^, and this can influence the net orientation of the solvent dipoles and the alignment of excess charges at the surface of condensates. Therefore, the presence of a surface, which drives the change of solvation environment ^23,31^, can also enable the formation of an electrically active interface. Further, the emergence of an electric field at a surface is governed by various mechanisms, suggesting its widespread occurrence ^61,142^. These factors might also support the idea that the catalytic activities should be generalizable for macromolecular structures ^143^. Further, the microenvironments within the dense phase encode unique ionic environment, micropolarity and water activities ^17,22,144,145^, thereby modulating various factors that can be critical to the chemical activity of condensates. Lastly, beyond microscopic structures, phase transition can also mediate the formation of subsaturated clusters in the dilute phase^73,74^. This might possess similar properties as micelles, which can also drive spontaneous catalytic activities^146^. Understanding distinct phase transition processes should further enhance our knowledge on the functional landscape of biomolecular structures.

### A role for water

Besides the asymmetric distribution of water molecules, we infer, based on our experimental characterizations, that the hydrogen bonding of water is heterogeneous in the condensates. More free water molecules exist within the condensates, especially at the interfaces of condensates, which supports our hypothesis regarding the roles of interfaces on modulating the catalytic functions. The distinct water activity can be caused by differences in the interfacial electric field or by variations in surface residues, which may have distinct affinities for water molecules or ions, thereby influencing the water structure. These observations emphasize the idea that the condensation not only alters molecular distribution but also molecular activity. Considering the diverse chemical reactions that water activity and structure play a critical role ^147,148^, understanding the molecular grammars regulating water will shed light on the chemical functions of Condenzymes.

### Can Condenzymes encode substrate-specificity?

Though the functions are encoded in mesoscale electrochemical environments, condensates possess distinct interior and interfacial structures based on their molecular compositions ^66,141^. These features can potentially drive specific residues onto the interfaces of condensates, similar to the defined surface residues on amyloid assemblies ^143,149^. These residues can encode binding selectivity toward specific chemicals, thereby modulating the specificity of catalytic functions ^150^. We expect that with enhanced understandings on the molecular structures of condensates ^151^, condensate-specific chemical functions will be defined.

Our study uncovers diverse factors that can affect the inherent catalytic behaviors of condensates. This does not require that condensates be formed by molecules with an intrinsic enzymatic functionality. Instead, just the formation of condensates via phase separation is sufficient, providing their unique electrochemical properties in the dense phase and the interface. Given that condensates are defined by their unique interphase potentials that arise from differences in chemical environments across coexisting phases and the ubiquitous existence of condensates ^152^, our discovery of inherent chemical functions of condensates suggests a possible role of condensates in maintaining cellular chemical homeostasis and managing cellular resources. This leads us to the possibility that condensates have the potential to be condenzymes.

The functions of condensates have remained elusive, and this has opened the door to criticisms and suggestions that condensates might simply be incidental bystanders in cells ^116^. Our findings appear to change this picture, with the upshot being that cells must have evolved mechanisms to leverage the inherent catalytic functions of condenzymes or cells have adapted mechanisms that blunt condenzyme functions. If the former is true, then working out condenzyme functions in modern cells becomes of utmost important. If the latter is true, then gains-of-function through condenzyme functions of pathological condensates becomes a challenge for cells, thus opening a new way to think about disease. Either way, our work opens a new way of thinking about condensates. It adds to recent discoveries of condensates having distinct and tunable electrochemical and viscoelastic properties ^11,12,14,22,23,30,98,153–158^. While the former discovery shows that condensates can be defined by membrane-like potentials that affect electrochemical equilibria in cells ^22,23,98^, the latter has broad implications for condensates being mechano-sensitive and mechano-responsive elements ^156,159^. The current work adds a biochemical dimension to the electrochemical and viscoelastic considerations of condensate functions. Indeed, our work, integrated with contributions made recently ^11,12,14,22,23,30,98,153–159^, opens the door to thinking about condensates being “coupled electro-chemo-viscoelastic” elements ^160^.

## RESOURCE AVAILABILITY

### Lead contact

Please direct requests for resources and reagents to Yifan Dai.

### Material availability

Plasmids generated in this study are available upon request to Yifan Dai.

### Data and code availability

- Raw data as part of this work is publicly available via the Dai lab Github and will be released to zenodo upon publication.
- All original code that was developed as part of this work is deposited to the Pappu lab Github repository is publicly available.
- Any additional information is available from the lead contact upon request.

## ACKNOWLEDGMENTS

This work was supported by the James J. McKelvey school of engineering (Y.D.) and the Center for Biomolecular Condensates at Washington University in St. Louis (Y.D., and R.V.P), the US Air Force Office of Scientific Research (FA9550-20-1-0241 to R.V.P, FA9550-21-1-0170 to W.M., and R.N.Z.), the St. Jude Collaborative on the Biology and Biophysics of RNP granules (to R.V.P), the US National Science Foundation (MCB 2419680 to R.V.P), and the US National Institutes of Health (R35 GM149256 to W.M., and F32GM146418-01A1 to M.R.K).

## AUTHOR CONTRIBUTIONS

Conceptualization: Y.D.

Methodology: Y.D., R.V.P., W.M., M.W.C., N.Q., R.N.Z., J.L., M.F., X.G.

Investigation: M.W.C., X.G., N.Q., X.S., M.F., Y.X., A.N., L.Y., Y.M., W.Y., M.R.K., V.L.

Funding acquisition: Y.D., R.V.P., W.M., R.N.Z., J.L.

Supervision: Y.D., R.V.P., W.M., R.N.Z., J.L.

Writing: Y.D., R.V.P., M.W.C., N.Q.

Reviewing & editing: all authors.

---

**Figure S1.**
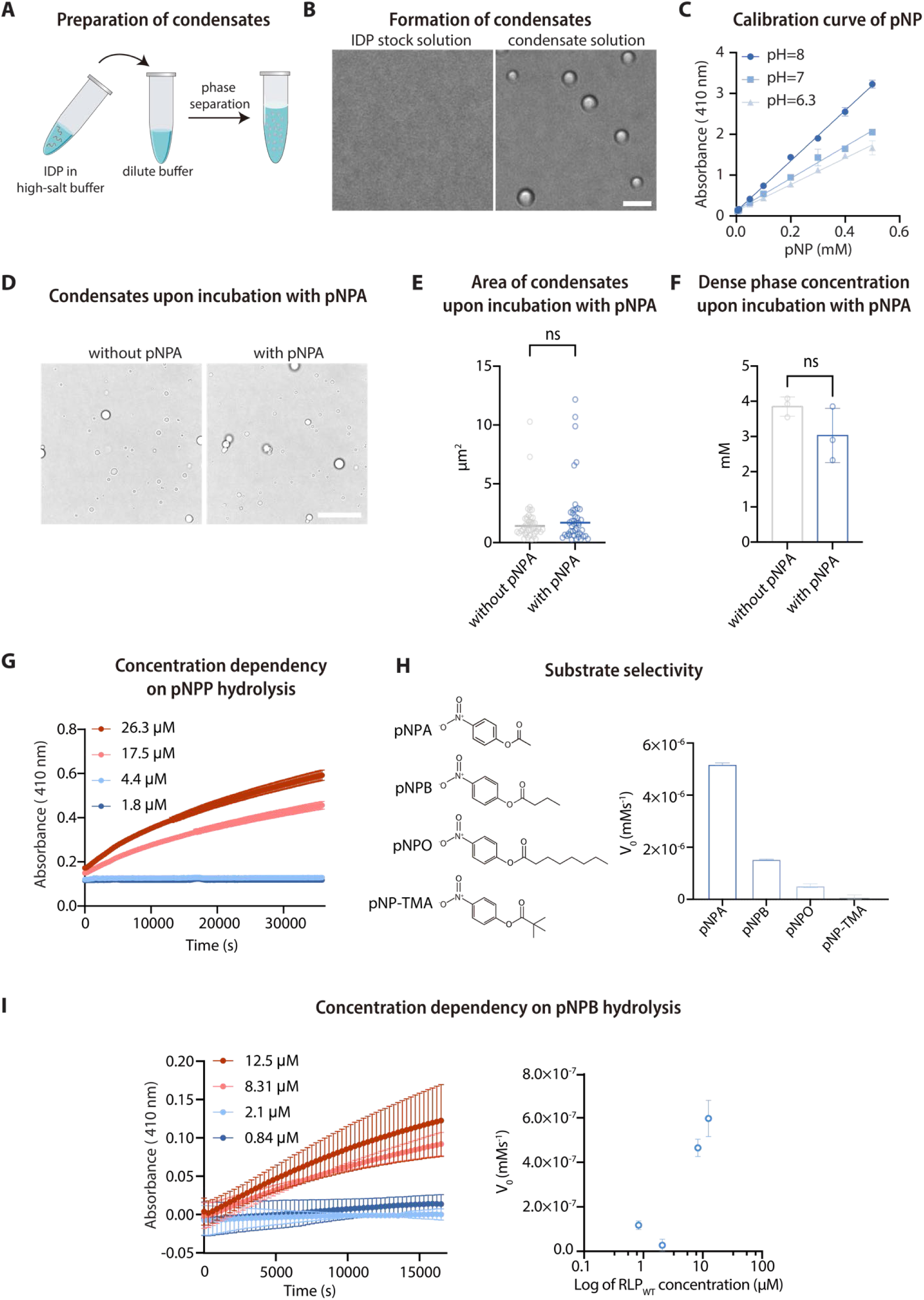
Evaluation of the hydrolytic capability of condensates with p-nitrophenol derived probes. **A,** Purified IDP is stored in a high-salt buffer, in which the protein remains soluble. The protein stock solution is added into a dilution buffer to trigger phase separation and the solution is incubated at room temperature for 30 min to allow condensate formation. For catalytic reaction, a high-concentration substrate stock solution is added into the condensate solution in a volume ratio of 1:200 to minimize the effects of the substrate solvent (e.g., acetonitrile) on condensate stability. **B,** Representative phase contrast images of IDP stock solution and condensate solution. Scale bar, 5 μm. **C,** Calibration curves of the concentration of the reaction product (p-nitrophenol) and the optical absorbance at 410 nm was constructed based on different buffer conditions. These buffer conditions cover the testing conditions in this study. **D,** Bright-field confocal images of RLP_WT_ condensate samples with or without the addition of pNPA incubated for the same amount of time (120 min). For the sample without the addition of pNPA, same volume of acetonitrile was added, which is the solvent of the pNPA stock solution. Scale bar, 20 μm. **E,** Quantification of the areas of individual condensates resided on the cover glass. ns, non-significant based on unpaired t-test with p=0.1057. N=50. **F,** Sedimentation assay quantification of the dense phase concentration of the RLP_WT_ condensate samples with or without the addition of pNPA. The condensate solution was dialyzed against the same reaction buffer without pNPA to remove the reactants and the products before subjecting the samples to sedimentation assay. ns, non-significant based on unpaired t-test with p=0.1587. N=3. **G,** Raw data curves for the analysis of the concentration-dependent hydrolytic capability of RLP condensates towards pNPP. N=3 **H,** Evaluation of the catalytic specificity of RLP_WT_ condensates using different substrates by comparing their initial rate at the same substrate concentration of 0.5 mM. The evaluated substrates were p-nitrophenyl butyrate (pNPB), p-nitrophenyl octanoate (pNPO) and p-nitrophenyl trimethylacetate (pNP-TMA). Chemical structures of pNPA, pNPB, pNPO, and pNP-TMA are displayed on the left. **I,** Raw data curves for the analysis of the concentration-dependent hydrolytic capability of RLP condensates towards pNPB, and the calculated kinetic values from these curves. N=3

**Figure S2.**
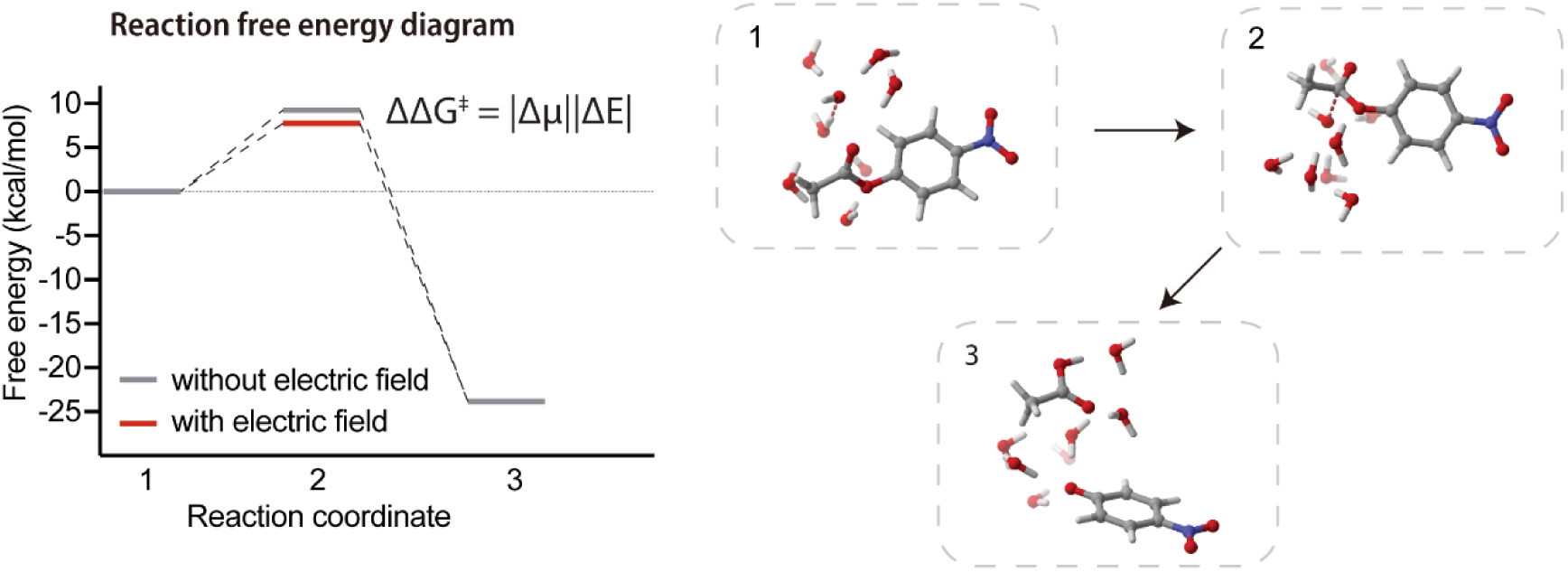
Evaluation of the role of interfacial electric fields on driving hydrolysis reactions. Reaction free energy diagram calculated based on DFT shows the effect of external electric fields on lowering the free energy barrier of the transition state of pNPA hydrolysis. The right panel shows the entire hydrolysis reaction pathway calculated with an explicit first solvation shell (with 7 H_2_O molecules) combined with a conductor-like polarizable continuum model (CPCM).

**Figure S3.**
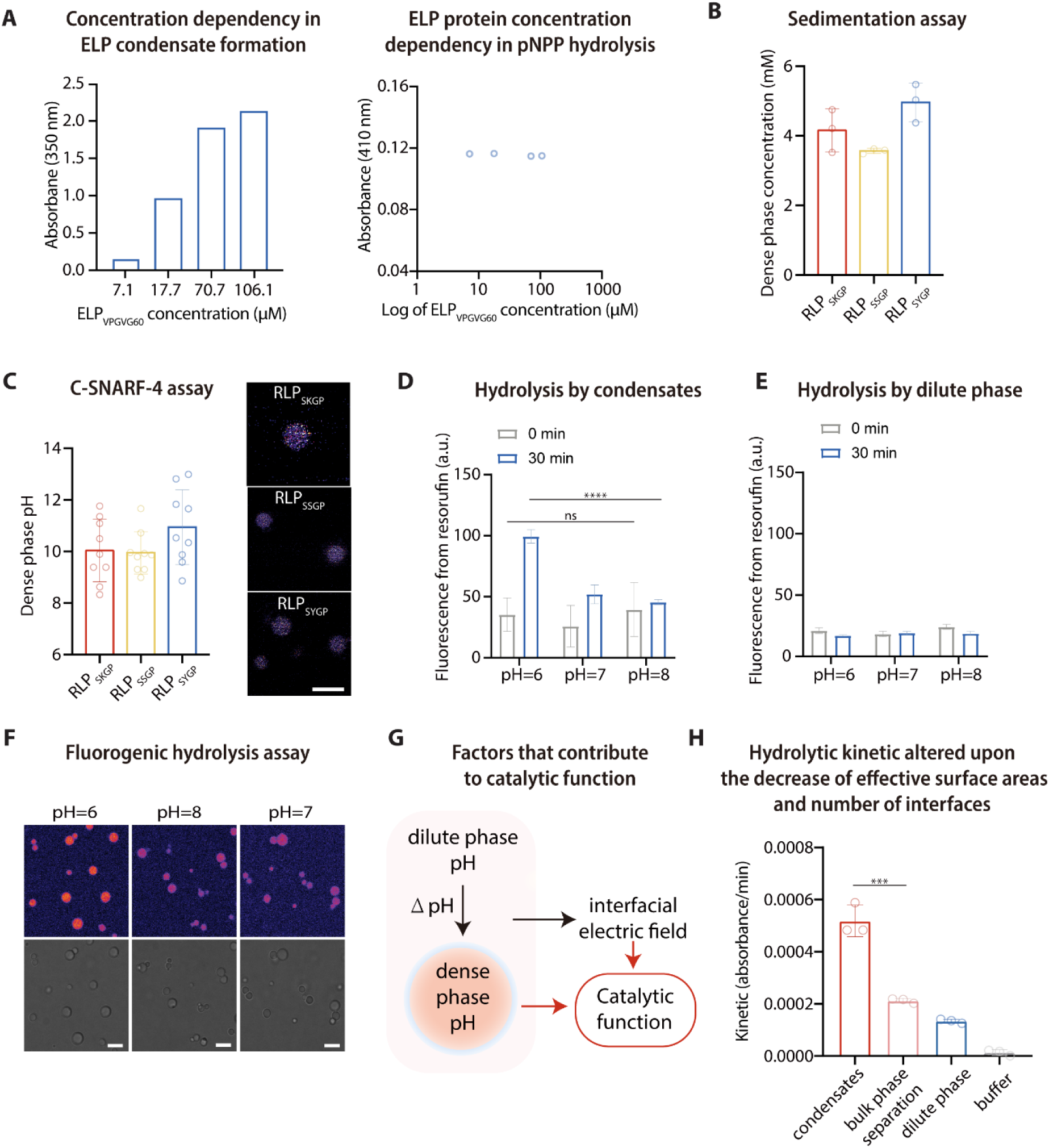
Evaluation of the contribution of dense phase, interfacial electric field, and interface on the catalytic capability of condensates. **A,** ELP protein concentration-dependent initial rate (V_0_) of the pNPB catalytic reaction. Similar to RLP proteins, an increase in ELP concentration leads to condensate formation, as evidenced by increased turbidity. However, unlike RLP proteins, increasing the ELP protein concentration does not lead to a sharp increase in the pNPP hydrolysis rate, suggesting that neither ELP proteins nor ELP condensates possess hydrolytic capability. **B,** Condensates were incubated for 2 h at room temperature before conducting the following characterizations. Sedimentation assay for the evaluation of the dense phase concentration of condensates formed by RLP sequences containing different N-terminus sequences. Compared between sequences, a slightly higher dense phase concentration was observed in the condensates formed by RLP_SYGP_. This observation is explained by the mutation from Lys to Tyr in the N-terminus sequence, which should mediate a stronger dense phase interactions. N=3 independent experiment. **C,** C-SNARF-4 assay analysis of the interior condensate pH formed by different sequences (RLP_SKGP_, RLP_SSGP_, RLP_SYGP_). No significant difference between each sequence was observed. This observation supports that the main sequence of RLP ([GRGDSPYS]_20_) determines the dense phase pH of condensates. N>8 global analysis of individual ratiometric images. Representative images from the C-SNARF-4 assay analysis of interior condensate pH formed by different sequences are displaying on the right. Scale bar, 5 μm. **D,** Quantified results from resorufin phosphocholine-based assay analysis of the hydrolytic capability of RLP_SKGP_ condensates at different pH. **E,** Quantified results from resorufin phosphocholine-based assay analysis of the hydrolytic capability of the dilute phase of RLP_SKGP_ condensates at different pH. **F,** Representative images from resorufin phosphocholine-based assay analysis of the hydrolytic capability of the dilute phase of RLP_SKGP_ condensates at different pH. The images were captured 30 min post reaction. Scale bar, 5 μm. **G,** Dominating factors contributing to the catalytic functions of condensates are the dense phase chemical environment and the interfacial electric field. **H,** pNPP probe-based hydrolysis assay analysis of the change of hydrolysis kinetic upon the decrease of effective surface areas and number of interfaces. N=3 independent experiments.

**Figure S4.**
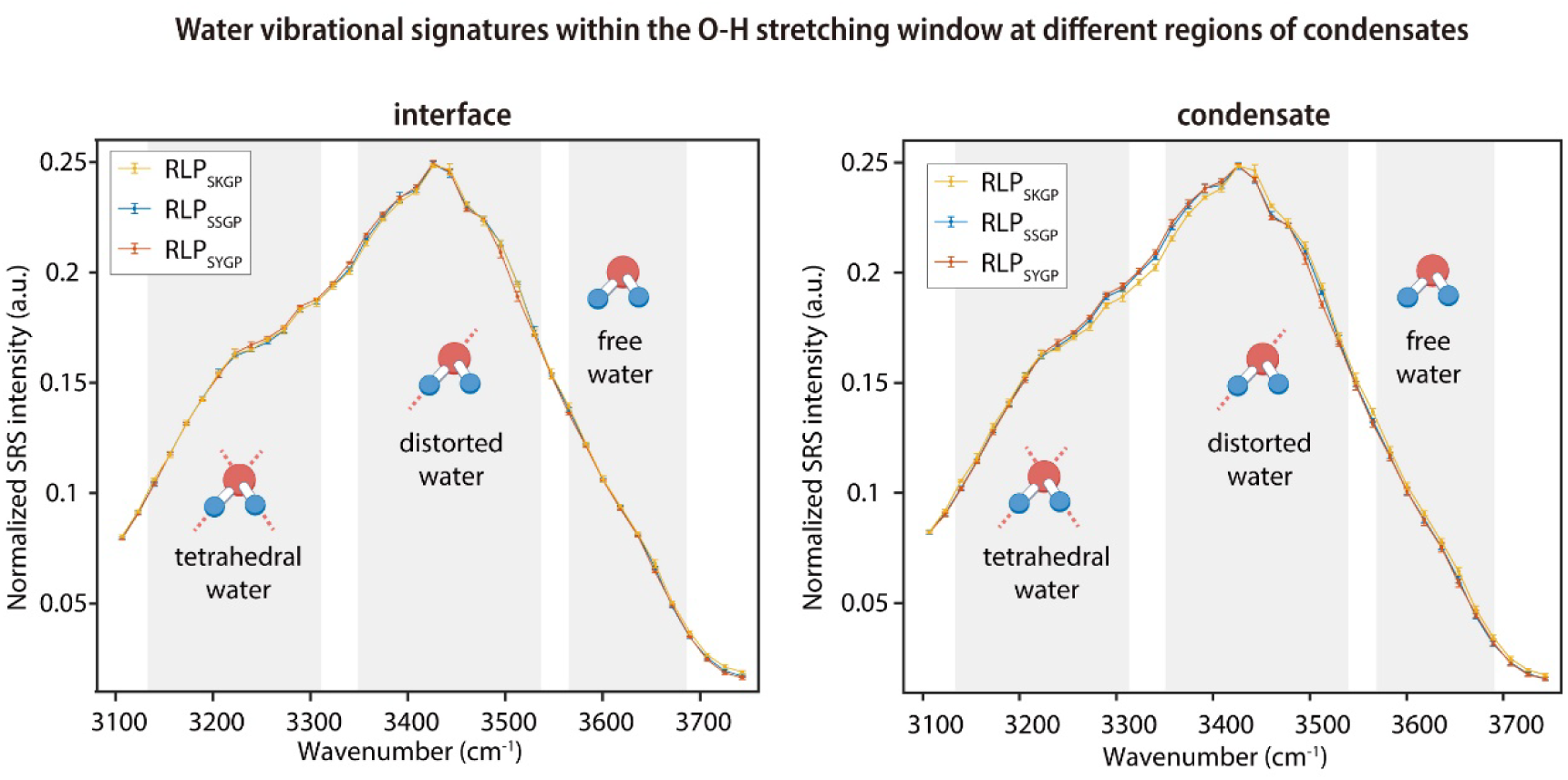
Stimulated Raman scattering (SRS) microscopy analysis of the water vibrational signatures of O-H bond at the interface and the interior region of condensates. This analysis was conducted on RLP protein with sequences, which are RLP_SKGP_, RLP_SSGP_, RLP_SYGP_.

**Figure S5.**
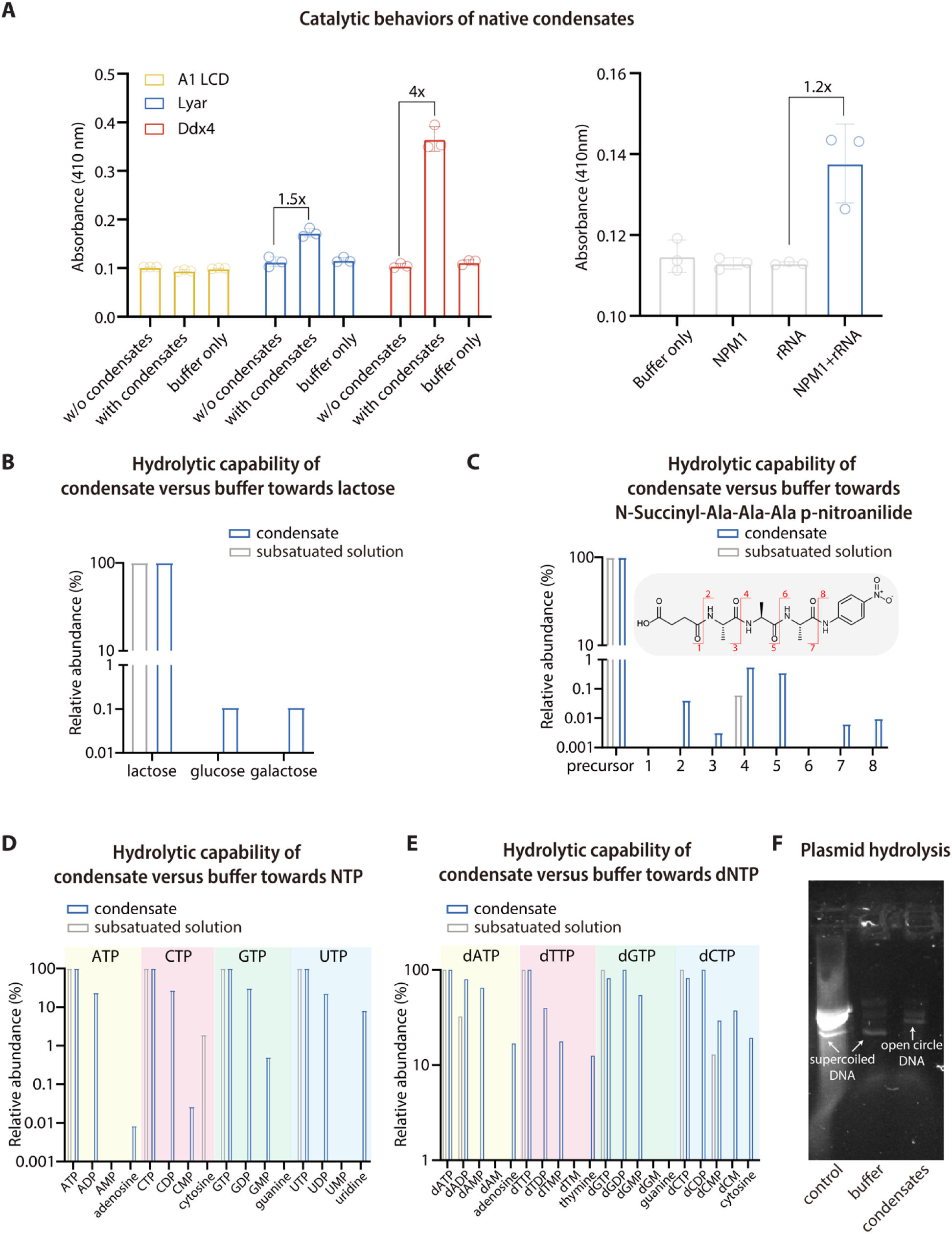
Evaluation of the general hydrolytic capability of native protein condensates and coacervates, and the general hydrolytic capability of the RLP_SKGP_ condensates towards different biomolecules. **A,** Evaluation of the hydrolysis of pNPP by different condensates formed by native IDPs and complex coacervation between NPM1 and rRNA. The hydrolysis reaction was conducted with 4 mM pNPP at 30 °C for 150 min. For condensates formed by IDPs only, the catalytic performance was compared between solutions containing IDP below Csat, condensates and buffer only. For condensates formed by complex coacervation between NPM1 and rRNA, the catalytic performance was compared between solutions containing NPM1, rRNA, condensates by NPM1 and rRNA and buffer only. N=3 independent experiments. **B,** Mass spectroscopy analysis of difference of the hydrolytic capability of condensates and protein solution without condensates (*c*_RLP_ < *c*_sat_) towards lactose. **C,** Mass spectrometry analysis of the differences in the hydrolytic capabilities between condensates and protein solution without condensates (*c*_RLP_ < *c*_sat_) toward N-Succinyl-Ala-Ala-Ala-p-nitroanilide. The chemical structure of N-Succinyl-Ala-Ala-Ala-p-nitroanilide is shown, with cleavage sites illustrated by red lines and eight fragments (four pairs) of hydrolysis products indicated by numbers. The precursor refers to N-Succinyl-Ala-Ala-Ala-p-nitroanilide. **D,** Mass spectroscopy analysis of difference of the hydrolytic capability of condensates and protein solution without condensates (*c*_RLP_ < *c*_sat_)towards NTPs. **E,** Mass spectroscopy analysis of difference of the hydrolytic capability of condensates and protein solution without condensates (*c*_RLP_ < *c*_sat_)towards dNTPs. **F,** DNA gel electrophoresis assay analysis of the changes in plasmid structure following incubation with either condensates or buffer revealed that the DNA plasmid lost its supercoiled conformation after incubation with condensates, resulting in an open circle structure. This alteration of plasmid structure indicated the condensates can cleavage the phosphodiester bond formed between the two DNA strands of the plasmid.

**Figure S6.**
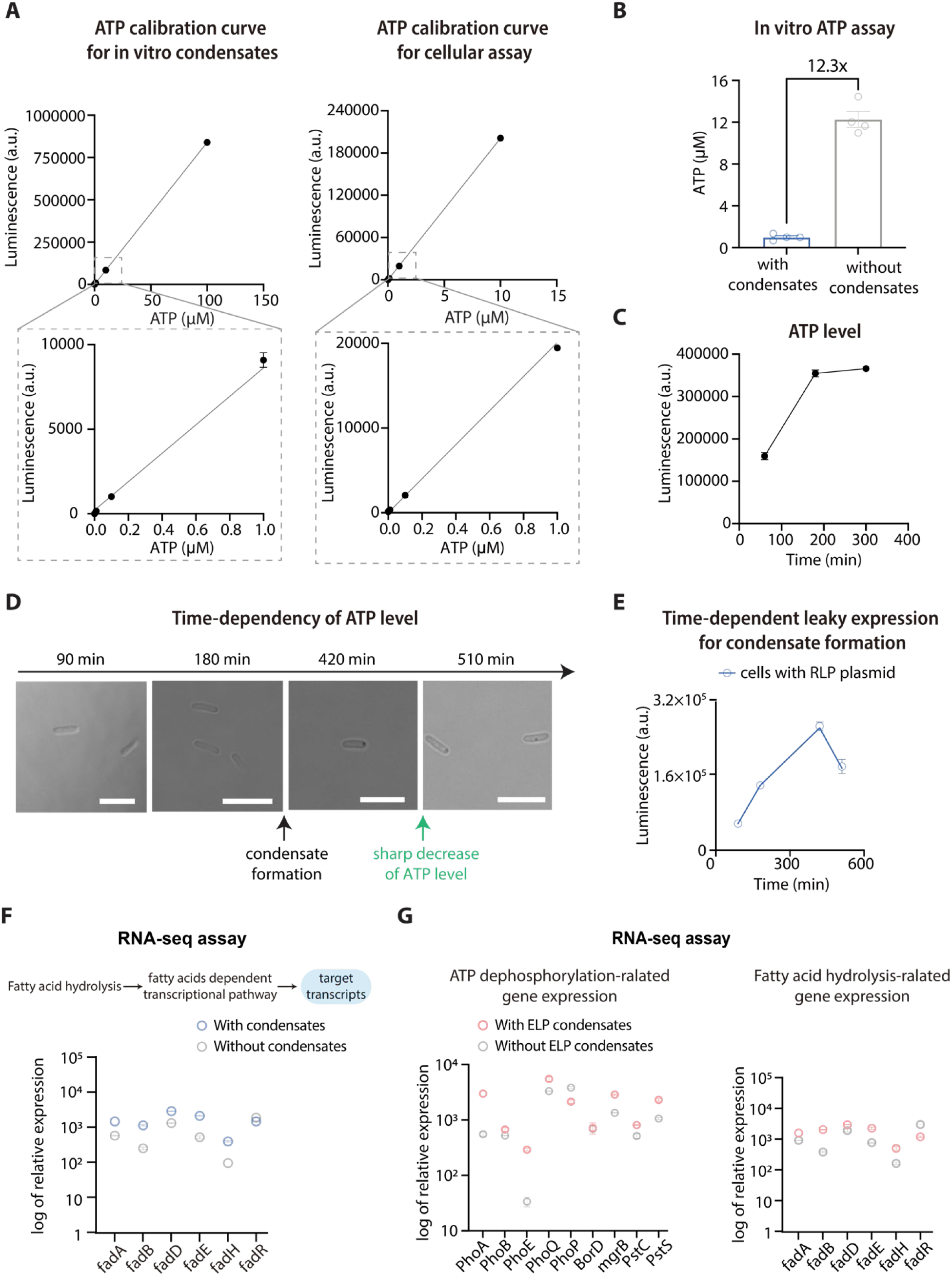
Evaluation of the ATP hydrolysis by condensates in vitro and in cells, and assessment of gene expression changes induced by the hydrolytic activity of condensates. **A,** ATP calibration curve constructed in the condensate formation buffer and the cellular assay buffer. **B,** Quantification of the decomposition of ATP based on solutions with or without condensates with an ATP-dependent luminescence assay. N = 4 independent experiment. Bar graph shows the mean ± SD. **C,** Evaluation of the effects of gene overexpression on the level of cellular ATP. The overexpression was induced by 0.5 mM IPTG at time zero. **D,** Confocal images of phase contrast images of cells containing the plasmid encoding RLP_SKGP_ regulated by a leaky T7 promoter. Without induction, condensate formation was observed at around 7 h, after which a sharp decrease of intracellular ATP level was observed as shown in Figure S6 E. Scale bar, 5 μm. **E,** Time-dependent tracking of the intracellular ATP level at different cellular stages. The decrease of the ATP level aligns with the time needed to form condensates. N = 4 independent experiment. Data point represents the mean ± SD. **F,** RNA-seq assay analysis of the difference in fatty acid hydrolysis-related gene expression in cells with or without RLP condensates. N=2 independent biological repeats. **G,** RNA-seq assay analysis of the difference in ATP dephosphorylation-related and fatty acid hydrolysis-related gene expression in cells with or without ELP condensates. N=2 independent biological repeats.

**Figure S7.**
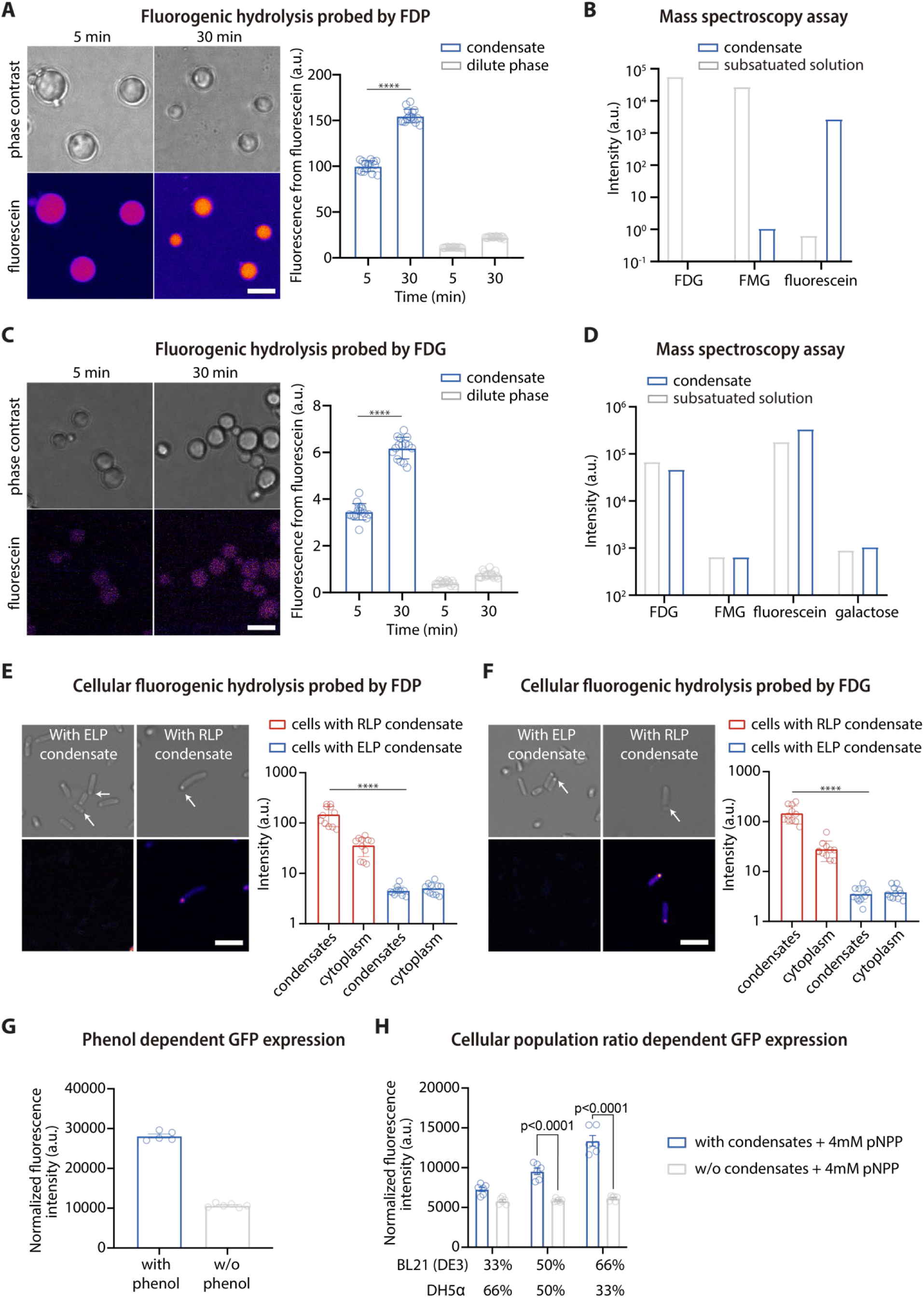
Verification of the hydrolytic capability of condensates in cells with fluorogenic hydrolysis probes and pNPP hydrolysis-based intercellular communication. **A,** FDP probe-based fluorogenic hydrolysis assay analysis of the hydrolytic capability of condensates. Data point represents the mean ± SD, with at least 11 data points. Scale bar, 5 μm. **B,** Mass spectroscopy assay analysis of difference of the hydrolytic capability of condensates and protein solution without condensates (*c*_RLP_ < *c*_sat_)towards FDP. **C,** FDG probe-based fluorogenic hydrolysis assay analysis of the hydrolytic capability of condensates. Data point represents the mean ± SD, with at least 11 data points. Scale bar, 5 μm. **D,** Mass spectroscopy assay analysis of difference of the hydrolytic capability of condensates and protein solution without condensates (*c*_RLP_ < *c*_sat_) towards FDG. **E,** Cellular fluorogenic hydrolysis assay analysis of the hydrolysis of FDP in cells with cells with RLP or ELP condensates. White arrows point to the condensates indicated by the phase contrast imaging. Data points represent the mean ± SD, with at least 15 data points. Scale bar, 5 μm. **F,** Cellular fluorogenic hydrolysis assay analysis of the hydrolysis of FDG in cells with RLP or ELP condensates. White arrows point to the condensates indicated by the phase contrast imaging. Data points represent the mean ± SD, with at least 15 data points. Scale bar, 5 μm. **G,** Comparison of the normalized GFP signal of cells with DmpR circuit with or without the addition of 100 μM of 4-nitrophenol after incubating for 24 h. **H,** Comparison of normalized GFP signal of a cell population containing different fractions of the BL21 (DE3) (with the condensate circuit) and DH5 (with the DmpR circuit) with the addition of 4 mM p-nitrophenol phosphate based on the conditions with or without condensates. P<0.0001 based on unpaired t-test. N=6.

## STAR METHODS

### Expression and purification of IDPs

#### Construction of RLP genes

The RLP_WT_ gene was a gift from Chilkoti lab^43,44^. The RLP_WT_ gene was modified through site-directed mutagenesis to generate mutants at the N-terminus using Q5 Site-Directed Mutagenesis kit (New England Biolabs). The cloned genes were transformed into NEB® 5-alpha Competent E. coli (High Efficiency) (New England Biolabs) and selected on a 2x YT plate with kanamycin. A single colony was picked and grown in 2x YT liquid medium (MilliporeSigma) containing 45 µg/mL Kanamycin at 37 °C for 18 h overnight (shaking at ∼225 r.p.m.). The cloned plasmids were then purified using QIAprep Spin Miniprep Kit (QIAGEN) and the DNA sequences of the purified plasmids were verified with Sanger sequencing service (Genewiz, Azenta Life Sciences).

#### Expression of synthetic IDPs

BL21 (DE3) competent *E. Coli* (New England Biolabs) was transformed with RLP genes (RLP_SKGP-WT_, RLP_SYGP-WT_, RLP_SSGP-WT_) or ELP genes (ELP_VPGVG-60_) and the transformed cells were selected by plating with Kanamycin. A single colony was picked and grown in 2 x YT liquid medium (MilliporeSigma) containing 45 µg/mL Kanamycin at 37 °C for 18 h overnight (shaking at ∼225 r.p.m.). The overnight cell culture was inoculated into 1 L 2 × YT liquid medium supplied with 45µg/mL Kanamycin. The bulk culture was first incubated at 37 °C (shaking at ∼225 r.p.m.) until the OD_600_ of cell culture reaching 0.5. The expression of IDPs was then induced by adding 0.5 mM final concentration of IPTG. For all the ELP proteins, the cell culture was incubated for another 18 h at 25 °C (shaking at ∼225 r.p.m.). For all RLP proteins, the cell culture was incubated for another 18 h at 37 °C (shaking at ∼225 r.p.m.).

#### Purification of RLP protein

Purification of RLP proteins was modified based on an established protocol using the upper critical solution temperature (UCST) feature of RLP^44,161^. Each liter of the overnight cell culture was pelleted by centrifugation at 2,000 g for 20 min at 4 °C (Eppendorf CR22N). Each pellet was resuspended in 30 mL Resuspension buffer (50 mM Tris, pH 7.5) and lysed by sonication (3 min total time, each cycle with 10 sec on and 40 sec off, 75 % amplitude) in an ice bucket. The cell lysate was centrifuged at 20,000 g for 25 min at 4 °C to separate the insoluble and the soluble phases. The insoluble pellet from each liter of bacterial culture was dissolved in 10 mL Urea buffer (50 mM Tris, 500 mM NaCl, 8 M Urea, pH 7.5) supplied with 500 units Benzonase Nuclease (MilliporeSigma) at room temperature for at least 2 h with rocking. The sample was then pre-warmed to 40 °C in a water bath for 30 min and centrifuged at 20,000 g for 25 min at 40 °C. The supernatant was collected and dialyzed into Resuspension buffer (50 mM Tris, pH 7.5). The dialysis buffer was changed one time over 3 h. Both the soluble and insoluble fractions in the dialysis bag were collected and centrifuged at 20,000 g for 25 min at 4 °C. The insoluble pellet was dissolved in Urea buffer (50 mM Tris, 500 mM NaCl, 8 M Urea, pH 7.5) at room temperature for at least 1 h with rocking. The sample was then warmed to 40 °C and centrifuged at 20,000 g for 25 min at 40 °C. The supernatant was collected, and its purity was determined on a SDS-PAGE gel (Bio-Rad, Any kD Mini-PROTEAN TGX Precast Protein Gels). This completes a round of UCST based purification. Another two rounds of temperature-dependent purification were performed before isolating the protein with AKTA size exclusion chromatography (cytiva). The final purity of the purified protein is higher than 95%.

The purified RLP protein was then processed in two different methods for storage. For liquid stock preparation, the concentration of the RLP stock protein was first adjusted to the concentration below its C_sat_ using Urea buffer and dialyzed into high salt buffer (50 mM Tris, 500 mM NaCl, pH7.5) at room temperature. The dialysis buffer was changed three times over 4 h. The dialyzed sample was first verified its capability to phase separate upon dilution. If not able to phase separate upon dilution, the concentration of the dialyzed sample was adjusted by the Pierce™ Protein Concentrators PES, 10K MWCO (ThermoFisher). Once it is verified, the protein stock was aliquoted into PCR tubes and frozen by liquid nitrogen and stored at −80 °C. For dry powder preparation, the rest of the purified RLP protein was dialyzed into milli-Q water at 4°C. The dialysis water was changed three times overnight. The dialyzed sample was then frozen at −80 °C for 1 h and lyophilized for three days to remove all water. The lyophilized dry powder can be stored at - 80 °C. We did not find a performance difference between samples stocked by different methods. The dry powder sample is more convenient for the preparation of RLP protein solution below C_sat_.

#### Purification of proteins for the generation of native condensates

##### Expression and purification of A1-LCD

BL21-CODONPlus RIPL *E. Coli.* cells (Agilent Technologies 230280) transfected with Δhexa_His-TEV-A1-LC 1 were grown in LB broth (Sigma: L3522) in Erlenmeyer flasks with ≥ 5-fold head volume at 37 °C (shaking at 220 r.p.m.) until OD600 ∼0.6 was reached. Cultures were then chilled for 15 min in an ice bath before adding 0.35 mM IPTG. Induced cultures were grown for additional 6 hours at 37 °C. Cells were harvested by centrifugation as described previously for synthetic IDPs, flash frozen, and stored as pellets in 50 mL Falcon tubes at −80 °C.

The cell pellet was resuspended to homogeneity in 35 mL supplemented lysis buffer (50 mM MES, 500 mM NaCl, 14.3 mM BME (β-Mercaptoethanol), 200 µM PMSF (phenylmethylsulfonyl fluoride) pH 6). Lysis buffer was then supplemented with: 500U DNAaseI (Sigma - 4536282001), 500U RNAse A (Sigma - 10109169001), 5 mg Lysozyme (Sigma - 62971-10G-F), and one protease inhibitor tablet (Sigma - 40694200). The cell suspension was lysed via sonication on a Branson 550 sonicator with an L102C horn attachment using five series of the following 20-round cycle: 1 second on / 2 second off at 30% power. The lysate was then pelleted through centrifugation for 30 min at 38,000 g. The lysate pellet was resuspended in a resuspension buffer (6 M GdmCl, 20 mM Tris, 15 mM imidazole, 14.3 mM BME, and 200 µM PMSF; pH 7.5) via repeat pipetting and pulses of sonication until the solution was visibly homogenous. This solubilized pellet was spun at 38,000g for 30 minutes at room temperature and the supernatant was applied to a gravity column with 5 mL bed volume of HisPur NiNTA resin (Thermo Fisher Scientific) equilibrated with resuspension buffer. The Ni-NTA resin was washed with 75 mL wash Buffer (20 mM Tris, 30mM imidazole, 4 M Urea, 14.3mM BME, and 200 µM PMSF; pH 7.5), then protein was eluted in elution buffer (20 mM Tris, 350mM imidazole, 4 M Urea, 14.3mM BME, and 200 µM PMSF; pH 7.5). Peak fractions from Ni-NTA affinity purification were pooled and diluted 1:1 in dilution buffer (20 mM Tris and 14.3 mM BME; pH 7.5). To this, 1 mg of TEV protease was added, and the mixture was dialyzed overnight in dialysis buffer (20 mM Tris, 2 M urea, 50 mM NaCl, 0.5 mM EDTA, and 1 mM DTT; pH 7.5) at room temperature. After 16 h, the protein solution was filtered through a 0.22 µm filter, then further purified via ion exchange chromatography on an ÄKTA Pure fast protein liquid chromatography (FPLC) module using a HiTrap SP 5mL column (Cytiva – 17115201). The column was equilibrated with buffer A (20 mM Tris, 2 M urea, 50 mM NaCl, and 14.3 mM BME; pH 7.5) and protein was bound then subjected to a contiguous gradient protocol with from 0.05M NaCl to 1 M NaCl. Fractions containing A1-LCD were pooled then further purified and buffer exchanged using size exclusion chromatography - HiLoad 16/600 Superdex 200pg column (Cytiva - 28989335) into the following storage buffer: 20mM MES and 4 M GdmCl at pH 5.5. Lastly, protein was pooled and concentrated in Amicon Ultra 3 MWCO (molecular weight cut-off) concentrator columns (Millipore-Sigma UFC 500396), according to the manufacturer’s suggestions. Concentrated protein was stored at 4°C for further usage.

##### Expression and purification of DDX4-NT

BL21 *E. coli*. cells (NEB) transfected with His-SUMO-DDX4-NT 2, a gift from Professor Lewis E. Kay, were grown in the same manner as A1-LCD. Cells were lysed in the same manner as A1-LCD with a different lysis buffer (20 mM Sodium Phosphate, 750 mM NaCl, 20 mM Imidazole, 14.3 mM BME, and 0.2 mM PMSF; pH 7.5). The supernatant was spun for 25 min at 38,000g and transferred into an equilibrated HisTrap FF Crude 5mL column (Cytiva – 11000458) with an ÄKTA Pure fast protein liquid chromatography (FPLC) module. The Ni-NTA column was washed in 75 mL lysis buffer then protein was eluted in elution buffer (0.02 M Sodium Phosphate, 0.5 M NaCl, 0.35 M Imidazole, 0.0143 M BME, 0.0002 M PMSF; pH 7.5). Peak fractions from this affinity purification were pooled and diluted 5-fold in dilution buffer (0.02 M Sodium Phosphate, 0.0143 M BME; pH 7.5). This solution was further purified via ion exchange chromatography using a continuous gradient purification protocol with a HiTrap Heparin HP 5mL column (Cytiva – 17040703), Buffer A (0.02M Sodium Phosphate, 0.1M NaCl, 0.0143M BME; pH 7.5), and Buffer B (0.02 M Sodium Phosphate, 0.1 M NaCl, 0.0143 M BME; pH 7.5) on the ÄKTA Pure FPLC module. Peak fractions containing SUMO-DDX4 protein were pooled and cleaved of SUMO tags during an overnight dialysis in the presence of 0.02x ULP1 Protease in cleavage buffer (0.02 M Sodium Phosphate, 0.3 M NaCl, 0.001M DTT (Dithiothreitol), pH 7.5). DDX4-NT was purified to ≥98% using size exclusion chromatography - HiLoad 16/600 Superdex 200pg column (Cytiva - 28989335) on the ÄKTA Pure FPLC module in storage buffer (0.022M Sodium Phosphate, 1.1M NaCl, 0.0143M BME, pH 7.5). The DDX4-NT solution was supplemented with 10% glycerol and concentrated in Amicon Ultra 3 MWCO (molecular weight cut-off) concentrator columns (Millipore-Sigma UFC 500396) and concentrated protein was aliquoted into single-use volumes (typically 10 µL), flash froze in liquid nitrogen, and stored at −80 °C. Each step of the purification was assessed in the same manner as described for A1-LCD.

##### Expression and purification of LYAR

Expression was carried out in the same way as described for A1-LCD. Cells were lysed in supplemented lysis buffer (20mM Sodium Phosphate, 500mM NaCl, 20mM Imidazole, 14.3mM BME, 200µM PMSF, pH 7). Supernatant was recovered from a 25-minute spin at 38,000g and transfered into a 10 mL volume of equilibrated HisPur Ni-NTA resin (Fisher - 88222) in a gravity column. The Ni-NTA column was washed in lysis buffer until no contaminant protein was detected by Bradford assay. Protein eluted in 20 mL elution buffer (20 mM Sodium Phosphate, 500 mM NaCl, 350 mM Imidazole, 14.3 mM BME, 200 µM PMSF; pH 7). Peak fractions from this affinity purification were pooled and diluted five-fold in dilution buffer (20 mM Sodium Phosphate, 14.3 mM BME; pH 7). This solution was further purified via ion exchange chromatography using a contiguous gradient purification protocol with a His-Trap Heparin HP 5mL column (Cytiva – 17040703), Buffer A (20 mM Sodium Phosphate, 100 mM NaCl, 14.3 mM BME; pH 7), and Buffer B (20 mM Sodium Phosphate, 100 mM NaCl, 14.3 mM BME; pH 7) on the ÄKTA Pure FPLC module. Peak fractions containing His-LYAR were pooled and purified to ≥95% using size exclusion chromatography - HiLoad 16/600 Superdex 75pg column (Cytiva - 28989333) on the ÄKTA Pure FPLC module in storage buffer (22mM Sodium Phosphate, 1.1M NaCl, 14.3mM BME, pH 7.5). The LYAR solution was supplemented with 10% (v/v) glycerol and concentrated in Amicon Ultra 3 MWCO (molecular weight cut-off) concentrator columns (Millipore-Sigma UFC 500396). Concentrated protein was aliquoted into single use volumes (typically 5 µL), flash frozen in liquid nitrogen and stored at −80 °C.

##### Expression and purification of NPM1

Expression was carried out in the same way as described for DDX4-NT and LYAR. Cells were lysed in supplemented lysis buffer (20 mM sodium phosphate, 500 mM NaCl, 20 mM imidazole, 14.3 mM BME, 200 µM PMSF, pH 7). Supernatant was recovered from a 25-minute centrifuge at 38,000 g and bound to a 10 mL volume of equilibrated HisPur Ni-NTA resin (Fisher - 88222) in a gravity column. The Ni-NTA column was washed in lysis buffer until no contaminant protein was detected by a Bradford assay. Protein eluted in 20 mL elution buffer (20 mM sodium phosphate, 500 mM NaCl, 350 mM imidazole, 14.3 mM BME, 200 µM PMSF; pH 7). Peak fractions from this affinity purification were pooled and diluted five-fold in dilution buffer (20 mM sodium phosphate, 14.3 mM BME; pH 7). Peak fractions containing His-MBP-Tev-NPM1 were pooled and cleaved of MBP during an overnight dialysis in the presence of 0.025x TEV protease in cleavage buffer (20 mM Sodium phosphate, 300 mM NaCl, 1 mM DTT, pH 7.5). This solution was further purified via ion exchange chromatography using a continuous gradient purification protocol with a HiTrap SP 5 mL column (Cytiva – 17115201), Buffer A (20mM sodium phosphate, 100mM NaCl, 14.3mM BME; pH 7), and Buffer B (20mM sodium phosphate, 100mM NaCl, 14.3mM BME; pH 7) on the ÄKTA Pure FPLC module. Peak fractions containing NPM1 were pooled and purified to ≥95% using size exclusion chromatography - HiLoad 16/600 Superdex 75pg column (Cytiva - 28989333) on the ÄKTA Pure FPLC module in storage buffer (22 mM sodium phosphate, 1.1 M NaCl, 14.3 mM BME, pH 7.5). The NPM1 solution was supplemented with 10% (v/v) glycerol and concentrated in Amicon concentrators. Concentrated protein was aliquoted into single use volumes (typically 5 µL), flash frozen in liquid nitrogen and stored at −80 °C.

##### Purification of Mature rRNA (mat-rRNA)

Mature rRNA (mat-rRNA) is a mixture of rRNAs (18S and 28S) and was purified using the same protocol described previously^11^. Briefly, rRNA was first purified from 50 µL packed volume of stage VI Xenopus oocytes Xenopus oocytes via TRIzol reagent extraction (Fisher - 15596026). A pellet of precipitated rRNA was reconstituted in 100 µL 4 °C ddH2O. Gel electrophoresis was carried out to confirm that the dominate RNA species were the 18s and 28s rRNA. rRNA Concentration was measured by absorbance at 260 nm on a Nanodrop 2000 and immediately aliquoted into single use volumes (typically 5 µL), flash-frozen in liquid nitrogen and stored at −80 °C.

##### Condensate formation for LYAR and DDX4

To prepare LYAR and DDX4 condensates, we mixed 3 µL of 2320 µM DDX4 (in 8 mM NaH_2_PO_4_, 12 mM Na_2_HPO_4_, 1 M NaCl) with 27 µL low-salt solution (8 mM NaH_2_PO_4_, 12 mM Na_2_HPO_4_, 0 M NaCl) into a 1.5 mL microcentrifuge tube and incubated for 30 min at room temperature before subjecting to catalytic assay. The corresponding buffer was used as base line to assess the spontaneous hydrolysis activity of the substrate in this buffer.

##### Condensate formation for A1 LC

To prepare A1 LCD condensates^68^, we mixed 27 µL of 100 µM A1-LCD (in 20 mM HEPES buffer, pH 7.0) with 3 µL of salt buffer (20 mM HEPES buffer with 3 M NaCl, pH 7.0) into a 1.5 mL microcentrifuge tube with a final concentration of 90 µM A1-LCD in 20 mM HEPES buffer with 300 mM NaCl. The condensate solution was incubated for 30 min at room temperature before subjecting to catalytic assay. The corresponding buffer was used as base line to assess the spontaneous hydrolysis activity of the substrate in this buffer.

##### Condensate formation through complex coacervation of NPM1 and rRNA

To prepare NPM1+mat-rRNA condensates, we mixed a 0.1x volume of 200 µM NPM1 (in mM NaH_2_PO_4_, 12 mM Na_2_HPO_4_, 1 M NaCl) with a 0.9x volume of low-salt solution (8 mM NaH_2_PO_4_, 12 mM Na_2_HPO_4_, 0 M NaCl) containing mat-rRNA at 11.12ng/µL. The condensate solution was incubated for 30 min at room temperature before subjecting to catalytic assay. The corresponding buffer was used as base line to assess the spontaneous hydrolysis activity of the substrate in this buffer.

#### Purification of ELP protein

Purification of ELP protein was modified based on an established protocol using the lower critical solution temperature (LCST) feature of ELP^162^. Each liter of the ELP cell culture was pelleted by centrifugation (Eppendorf CR22N) at 2,000 g for 20 min at 4 °C. Each pellet from 1 liter of cell culture was resuspended into 35 mL 1×PBS and lysed by sonication (3 min total time, each cycle with 10 sec on and 40 sec off, 75 % amplitude) in an ice bucket. 1 tablet of cOmplete™ Protease Inhibitor Cocktail (Roche) and 500 units of Benzonase Nuclease (MilliporeSigma) were then added into the cell lysate and incubated at room temperature for 1 h. The cell lysate was centrifuged at 20,000 g for 25 min at 4 °C to separate the soluble and the insoluble phases. The supernatant, which contains the soluble ELP, was collected in a new tube. The ELP phase transition was then induced by adjusting the NaCl concentration to 1 M and heating in a 40 °C water bath for 20 min. The turbid ELP solution was then pelleted by centrifugation at 20,000 g for 25 min at 40 °C. The insoluble ELP pellet was physically disrupted using pipette tip and dissolved in 20 mL fresh 1×PBS at 4 °C for at least 3 h with rocking. The dissolved ELP was centrifuged at 20,000 g for 25 min at 4 °C and the supernatant was collected. This completed a full round of ELP purification based on its LCST phase behavior. Another two rounds of temperature-dependent purification were performed before isolating the protein with AKTA size exclusion chromatography (cytiva). Protein purity was then determined on an SDS-PAGE gel (Bio-Rad, Any kD Mini-PROTEAN TGX Precast Protein Gels). This process would lead to a purity over 95%. The purified ELP was dialyzed into milli-Q water overnight at 4 °C. The dialysis water was changed three times. The dialyzed sample was then frozen at −80 °C for 1 h and lyophilized for three days to remove all water. The lyophilized dry powder was stored at −80 °C.

#### Catalytic activity assay

To prepare RLP proteins for the catalytic activity assay, a frozen protein stock was freshly thawed to room temperature before dilution. For the condensates-forming condition, each reaction contained 60 µL protein stock and 140 µL Dilution buffer (50 mM Tris, pH was adjusted to meet the specific testing condition described in the manuscript). For the dilute-protein (no condensates formation) condition, each reaction contained 10 µL protein stock, 50 µL High Salt buffer (50 mM Tris, 500 mM NaCl, pH 7.5) and 140 µL Dilution buffer (50 mM Tris, pH was adjusted to meet the specific testing condition described in the manuscript). The well-mixed protein solutions were incubated at room temperature for 30 min to allow condensates formation before adding substrates.

To prepare ELP condensates for the catalytic activity assay, lyophilized ELP powder was weighted to prepare a stock solution in Dilution buffer (50 mM Tris, pH 7.5). After the powder was fully dissolved, NaCl concentration was adjusted using a high concentration NaCl stock solution (50 mM Tris, pH 7.5, 2 M NaCl) to realize a final salt concentration at 150 mM NaCl.

To prepare chemical stock solutions, a 100× chemical stock was freshly prepared before every experiment. 4-nitrophenyl phosphate (pNPP) was prepared in water. 4-nitrophenol (4-NP), 4-nitrophenyl acetate (pNPA),, 4-nitrophenyl trimethylacetate (pNP-TMA) were prepared in acetonitrile. 4-nitrophenol butyrate (pNPB) and 4-nitrophenyl octanoate were prepared in dimethyl sulfoxide (DMSO).

To initiate the catalytic reaction after protein incubation, 2 µL of a 100× chemical stock solution was added per 200 µL of diluted protein sample on a flat-bottom transparent 96-well plate (Corning, COSTAR). For control reaction without chemical, the same volume of corresponding solvent was added. The plate was sealed with a transparent film and immediately transferred to a plate reader for recording. The absorbance was measured at 410 nm and 385 nm with 25 flashes on a TECAN Infinite M200 plate reader or a TECAN Multimode plate reader (TECAN). The reaction was evaluated at 25 °C with a 3-min kinetic interval for at least 70 cycles.

From the measured absorbance at 410 nm for all catalytic reactions, the concentration of 4-Nitrophenol, the product of the hydrolysis reactions, was calculated based on a standard curve which was obtained experimentally under the same condition. The baseline of 4-nitrophenol from chemical self-hydrolysis was subtracted from the catalytic reaction. The initial rate (V_0_) was calculated from the linear portion of the activity plot ([4-Nitrophenol] versus time) by performing a linear regression using Graphpad. For Michaelis-Menten fitting, the means of V_0_ values were plotted as a function of substrate concentrations, and the plot was fitted to the Michaelis-Menten equation by using the built-in non-linear regression on Prism Graphpad. The fitted Michaelis Menten equation was Y=V_max_*X/(K_m_+X). All measurements were replicated at least 4 times.

#### Absorbance scan

The absorbance scan of 0.8 mM pNPA hydrolysis reaction catalyzed by RLP_SKGP-WT_ condensates was conducted at 25 °C at pH 7.5. The reaction condition and the buffer composition were identical to the aforementioned catalytic activity assay. At different incubation time points, measurements were taken using the UV-VIS function on Nanodrop one (Thermo Scientific).

#### Fluorogenic hydrolysis assay

The condensate solutions were prepared as described in previous sections. Resorufin phosphocholine was added into the condensate solution at a final concentration of 50 μM. The reaction solution was transferred into a 384 well plate (PerkinElmer) and incubated in H301-K stage (okolab) at 30°C. The 10 % excitation was set at 550 nm with a WLL laser, and the emission detector was set at 570-600 nm on a HYD detector. An autofocus function was set to image a z range of 15 μm and the image with the highest fluorescence signal was used for quantification. ImageJ (Fiji) was used for quantification based on Particle Analysis function.

The fluorogenic hydrolysis assay was employed to probe the change of hydrolytic capability of RLP_SKGP_ condensates upon the change of dilute phase pH. In this experiment, the condensates were prepared as described in the previous section with the pH of the dilution buffer adjusted to 6, 7, and 8, respectively. After the formation of the condensates, the fluorogenic hydrolysis assay was implemented following the aforementioned protocol.

#### In vitro ATP dephosphorylation assay

To evaluate the level of dephosphorylation of ATP by RLP_SKGP-WT_ condensates *in vitro*, protein samples were prepared by the same method as in the aforementioned catalytic activity assay. The ATP stock solution (prepared from adenosine 5’-triphosphate disodium salt hydrate) was added to initiate the reaction with a final concentration of 10 µM. The reaction was incubated at 30 °C for 30 min or 90 min. The ATP quantification was performed by using the Luminescent ATP Detection Assay Kit (Abcam) following the manufacturer’s protocol. In brief, the reaction was terminated by adding the detergent and incubated for 5 min. The substrate solution was then added into the sample and incubated for 15 min in the dark. The luminescence was recorded on the TECAN Infinite M200 plate reader with a 2000 ms integration time (Luminescence mode). For the ATP calibration standard, the ATP standard solutions were freshly prepared for each assay under the same experimental condition.

#### *E. Coli.* intracellular ATP assay

To measure the level of intracellular ATP upon the formation of RLP_SKGP-WT_ condensates, the BL21 (DE3) competent *E. coli* was transformed with the RLP_SKGP-WT_ plasmid and the transformed cells were selected on a Kanamycin plate. A single colony of the plated BL21 *E. coli* was picked and grown in 2 mL 2× YT liquid medium supplied with 4 % glucose and 45 µg/mL Kanamycin at 37 °C overnight. Each liquid culture was back diluted at a ratio of 1:100 with fresh 2× YT liquid medium supplied with 45 µg/mL Kanamycin. After a 1.5-hour incubation at 37 °C, the culture was either induced with Isopropyl β-d-1-thiogalactopyranoside (IPTG) to produce RLP_SKGP-WT_ protein or not induced. At post-induction time points of 3 and 7 h, OD_600_ of each sample was measured and adjusted to the same cellular density using 1×PBS after washing with 1×PBS. The quantification of ATP level was conducted using the Luminescent ATP Detection Assay Kit (Abcam) following the manufacturer’s protocol. In brief, the reaction was terminated by adding the detergent and the sample was shaken in an orbital shaker for 5 min. The substrate solution was then added, and the sample was transferred to the TECAN Multimode plate reader for a 3-minute shaking, followed by a 12-minute dark adaption. The luminescence was recorded with 2000 ms integration time using the Luminescence mode.

#### Hydrolytic product detection

The nano-electrospray ionization mass spectrometry (nESI-MS) was employed for the target hydrolytic products detection. The nESI emitter was made of borosilicate glass capillary (Lot: 213425, ITEM# BT-150-10, ID: 0.86 mm, OD: 1.5 mm, Length: 10 cm) and fabricated by a micropipette puller (Sutter Instrument Co., Novato, CA, USA). Major parameters of the pull program were set as follows: HEAT 715; VEL: 15; PULL: blank; TIME: 250; PRESSURE: 720. Sample solution was aspirated and loaded by a micro-loader (Lot# K2080231, 20 μL, Eppendorf, Germany) into the fabricated capillary emitter. A stainless stain needle was inserted from the back side of the emitter to touch the loaded sample solution. The sample-loaded nESI emitter was then positioned in front of a mass spectrometer (Obitrap Fusion, Thermo Scientific, San Jose, CA, US) inlet at a distance of 3.0 mm. When a +1.5 kV high voltage is applied to the needle, the strong electric field form on the emitter tip will trigger the spraying process for the product compound ionization and MS detection. The target metabolite and its hydrolytic products were monitored by the full scan under a positive mode (m/z 100-1000). The MS capillary temperature was set at 275 °C; RF-lens (%) was set at 40, and maximum injection time set at 400 ms. The mass spectrometer was calibrated before each measurement batch by using ion mass calibration solution kits (Pierce positive, Thermo Fisher, San Jose, CA, USA). Before being detected by mass spectrometry, samples were incubated for 4 h at room temperature.

##### Mass spectroscopy assay analysis of the hydrolytic capability of condensates in cells

To conduct this experiment, cells were first processed so that we can obtain the intracellular metabolites. First, different groups of cells with an equivalent density level were harvested by centrifugation at 3500 rpm for 10 min at 4 °C. Thereafter, a flash frozen was conducted by using liquid nitrogen to quench all biochemical processes within cells. The cell samples were frozen under −80 ℃ until use. Before MS detection, acetonitrile will be added to the concentrated cell suspension to deactivate intracellular enzymes and make the final solvent composed of 90% acetonitrile-10% water in volume ratio. The diluted cell suspension will then immediately go through a 5 minutes’ ultrasonication (Branson 3800 ultrasonic cleaner, 40 KHz) to fully disrupt cell membranes and release intracellular metabolites into the solution. Finally, another 5000 rpm centrifugation will be conducted to spin down the insoluble cell debris and denatured proteins. Only the supernatant will be uploaded into a capillary emitter for mass spectrometry detection.

#### RNA-seq assay

To conduct the RNA-seq assay, BL21(DE3) was first transformed with plasmids containing RLP_SKGP-wt_, ELP_(VPGVG)60_, or the blank plasmid. These transformed cells were then plated on 2xYT agar with 50 mg/L kanamycin and 0.4% w/v glucose. Upon culturing, single colonies were inoculated into 3 mL of 2xYT supplemented with 50 mg/L kanamycin and 0.4% w/v glucose in 14 mL round bottom tubes (Fisher Scientific) and grown overnight in a shaking incubator at 37 °C and under constant shaking at 250 rpm. Then, cultures were 1% v/v back diluted into fresh M9 media until the exponential phase (OD600 = 0.2-0.25) was reached. 3 μL of 0.5M IPTG was added to the medium, and induced cells were continually cultured for 1 h. Finally, all cultures were centrifuged to obtain cell pellets, which were sent for RNA-seq analysis at Azenta Life Sciences.

#### Confocal microscopy for in vitro condensates

Analysis of condensate quantity and chemical environments were performed using LEICA STELLARIS 8 FALCON confocal microscopy (LEICA). For characterization of the chemical environments of the condensates, we followed the previous published protocols^14^ using SNARF™-4F 5-(and-6)-Carboxylic Acid (ThermoFisher) and DI-4-ANEPPS (Thermofisher) dyes for the characterizations of internal apparent pH and the relative strength of interfacial electric fields. The images were processed through ImageJ with Image Calculator to quantify the ratio of fluorescent signals between each channels. The calibration curve for pH is *y* = −0.1492 × *x* + 1.622 with *y* as the ratio of C-SNARF-4 signal and *x* as the pH condition.

#### Hyperspectral SRS imaging

Hyperspectral SRS imaging of condensates was acquired by a standard SRS microscope equipped with a picoEmerald FT laser system (Applied Physics & Electronics) to supply a synchronized pump beam and a Stokes beam with a 2 ps pulse width and 40 MHz repetition rate. The Stokes beam is fixed at 1031.2 nm and modulated at 20 MHz. The two beams are spatial-temporally overlapped before being coupled into an inverted laser-scanning microscope (FV1200, Olympus) and focused by x25 water objective (XLPlanN, 1.05 numerical aperture, MP, Olympus) for SRS imaging. The transmitted beams are then collected by the 1.4 NA oil condenser and filtered with a shortpass filter (FESH1000) to leave only the pump beam to be collected by a silicon photodiode (FDS1010, Thorlabs) with a DC voltage of 64 V. The current output from the photodiode was terminated by 50 Ω and demodulated with a high-frequency lock-in amplifier (HF2LI, Zurich Instruments) at 20 MHz frequency to detect the stimulated Raman loss on the pump beam. The in-phase X-output of the lock-in amplifier is fed back into the analog interface box of the microscope to form SRS images.

For mapping the vibrational signature of water OH stretching, the pump beam was tuned in the range between 741 nm and 802 nm. The hyperspectral images were acquired with a pump power of 17 mW and Stokes power of 125 mW on the sample. The time constant at the lock-in amplifier was set at 18 μs, with the pixel dwell time for scanning set to 20 μs.

#### LaSSI Simulations

To evaluate whether the Simulations were performed using LaSSI, a lattice-based Monte Carlo engine^65^. Monte Carlo moves are accepted or rejected based on the Metropolis–Hastings criterion so that the probability of accepting a move is equal to min (1, exp(–*β*Δ*E*)), where *β* = 1 / *kT*. Here, *kT* is the simulation temperature, and Δ*E* is the change in total system energy associated with the attempted move. Total system energies were calculated using a nearest neighbor model and previously derived interaction parameters^28^. For each variant and simulation temperature, 300 distinct chain molecules composed of 166 beads each were placed in a cubic lattice with a length of 140 lattice units. Previous calibrations have shown that the numbers of molecules used in these simulations are adequate to avoid problems due to finite size effects^28^. The simulations were allowed to equilibrate such that the chains formed a single condensate with a coexisting dilute phase before any analysis was performed. This process was facilitated by initializing the system in a smaller box with side length 40 lattice units. Simulations were performed for 3 × 10^10^ Monte Carlo steps and analyzed every 5 × 10^7^ steps. Radial densities were determined by calculating the center-of-mass of the system and partitioning the system into radial bins, centered at the center-of-mass, each with radius 1/4 of a lattice unit. Radial densities were normalized by the exact number of lattice sites within each radial bin. Radial densities were further scaled based on the number of beads being analyzed. For example, when determining radial densities for the full-length proteins, densities were divided by 166 (the chain length). In contrast, when determining radial densities for the N-termini, the first four residues of the protein were used, so the densities were divided by 4. For each set of conditions, three independent simulations were performed.

#### Density functional theory calculations

To understand the role of electric field on catalyzing the hydrolysis reactions, we investigated the influence of the external electric field strength on the rate of PNPA hydrolysis computationally using density functional theory (DFT). It has been shown computationally that PNPA hydrolysis is likely concerted with the tetrahedral intermediate being the transition state, and the reaction is well-studied experimentally^163,164^. In the presence of an electric field, the free energies is modulated by −*µ*·**E**, where *µ* is the dipole moment of any species and **E** is the electric field. The barrier height changes for this reaction due to the electric field is given by

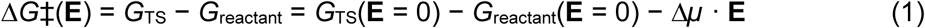

where Δ*µ* = *µ*_TS_ − *µ*_reactant_ is the difference in dipole between the reactant and the transition state. From this, the field-induced reaction barrier change is expressed as ΔΔ*G*^‡^ = −Δ*µ* · **E**. By employing the Arrhenius equation, the expected rate change was calculated through the equation,

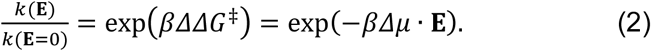

All computations were performed with Q-Chem 6.0^57^. The dense phase environment was modeled using the conductor-like polarizable continuum model (CPCM), a standard implicit solvation model to model these reaction systems^165,166^

We investigated a model with an explicit solvent cage^163^ along with implicit solvation treatment. We found that without adding an explicit solvent cage, the barrier height was too low (1−2 kcal/mol) for various exchange-correlation functionals and basis sets employed. Therefore, an explicit solvent cage consisting of seven water molecules was added around the reaction complex, following a published study^163^. To model the system, the 6-31++G** basis set and B3LYP exchange-correlation functional was first used to compute optimized gas-phase geometries for reactant, transition state, and product. The optimized gas-phase geometries with implicit solvation were studied via CPCM, the def2-TZVPD basis set, and *ω*B97X-V exchange-correlation function. The solvation-free energies and dipoles for the reaction system were obtained for the reactant, transition state, and product configurations. The dipoles for the reactant and transition state and change in barrier height due to a changing electric field were computed for several dielectric constants *ε* since we cannot measure the precise dielectric constant. Modulating the dielectric constant does not change the qualitative conclusions. We reported the free energy changes based on *ε* = 3.2, the average dielectric constant for proteins^167^ and the electric field of 10 MV/cm (a typical value in the protein environment^58^)

#### Isolation of the contribution of interface on the catalytic reactions by decreasing the effective surface areas and number of condensates

To conduct this experiment, the RLP_SKGP_ protein was first purified and diluted to an A280 of 3.0 in a storage buffer (4 M Urea, 500 mM NaCl, 50 mM Tris, pH 7.5). The protein was then dialyzed into a buffer with a significantly lower salt concentration (150 mM NaCl, 50 mM Tris, pH 7.5) to directly trigger the formation of condensates. The dialysis process lasted for 3 hours. The solution containing condensates was then collected for subsequent experiments.

A portion of the condensate-containing solution was centrifuged (20000g, 20 minutes) to pellet the condensates, thereby lowering the effective surface area and number of interfaces. Subsequently, the supernatants were collected and filtered through a membrane that allow the passage of molecules with a molecular weight lower than 100 kDa, thereby eliminating nano-sized protein assemblies. The solution after this filtering was designated as the dilute phase. Subsequently, half of the dilute phase solution was refilled into the tubes containing the protein condensate pellet. The refilling of the diluted phase was as conducted as gentle as possible while ensuring the fully contact with pellets. Finally, pNPP was added to the following solutions: the solution with condensates, the dilute phase solution refilled to the tubes with condensate pellets, the dilute phase itself, and the buffer from dialysis. All solutions were in the same volume, with a final pNPP concentration of 5 mM. The absorbance changes of these solutions at 410 nm were monitored using a plate reader at different time points.

#### Isolation of the contribution of interface on the catalytic reactions by adding charged surfactants

To conduct this experiment, RLP_SKGP_ condensate was prepared by diluting the protein stock in high salt buffer (500 mM NaCl, 50 mM Tris, and pH=7.5) with a Tris buffer (50 mM Tris, and pH=7.5) with a volume ratio of 3:7. Upon the formation of condensates, charged surfactants, SDS and CTAB, were supplement into the solution containing condensates with a final concentration of 100 μM, separately. The surfactants were then incubated with condensates for 10 min. Subsequently, ANEPPS assay was conducted to evaluate the effect of charged surfactants on the interfacial electric field of condensates. Further, fluorogenic hydrolysis assay based on the resorufin phosphocholine probe was employed to analysis the effect of charged surfactants on the hydrolytic capability of condensates.

#### Cellular fluorogenic assay

Two fluorogenic hydrolysis probes were employed in this experiment: FDP (fluorescein diphosphate) and FDG (fluorescein di(β-D-galactopyranoside)). Both FDP and FDG are well-established probes for analyzing cellular metabolism. The ability of condensates to hydrolyze both probes was first verified in vitro. In these experiments, FDP or FDG was added to a solution containing condensates to a final concentration of 1 mM. If either probe is hydrolyzed to generate fluorescein, the fluorescence intensity is expected to increase significantly. This change in fluorescence was monitored using confocal microscopy with an excitation wavelength of 490 nm and an emission range of 510–530 nm. Upon incubation with condensates, the fluorescence intensities in the samples increased, indicating that both probes were hydrolyzed by the condensates.

As for cellular hydrolysis assay, the hydrolysis of FDP and FDG probes are monitor upon incubation with cells with RLP_SKGP_ condensates, cells with RLP_SKGP_ plasmid but not yet formed condensate, and cells with ELP condensates. Before characterization, cells were dilute to a density with OD600=1. Subsequently, FDP and FDG probes were added into the medium with a final concentration of 100 μM. Both probes were incubated with cells formin. After the incubation, cells were imaged with STELLARIS 8 FALCON confocal microscopy (LEICA), followed by quantification with ImageJ (NIH).

#### Hydrolysis of DNA plasmid by RLP condensates

The DNA plasmid was incubated with solutions with RLP_SKGP_ condensates or buffer for 30 min at room temperature. Subsequently, proteinase K was added to the samples to degrade the RLP_SKGP_ protein for 30 min at 37 °C. The samples were then subjected to agarose gel electrophoresis to analyze changes in plasmid structure or molecular weight, compared to an untreated plasmid as a reference. Finally, the agarose gel was analyzed using the GelDoc Go imaging system (Bio-Rad).

#### Hydrolysis of ssDNA by RLP condensate

An ssDNA hydrolysis probe containing 6-FAM at the 5’ end and BHQ-1 at the 3’ end was purchased from IDT. Experimentally, the 6-FAM-ssDNA-BHQ-1 probe (40 nM) was incubated with a solution containing condensates. Subsequently, confocal microscopy was employed to monitor fluorescence emission from the sample at a wavelength of 510–530 nm with an excitation laser at 495 nm.

#### Inter-cellular communications with DmpR circuit

To evaluate whether the catalytic activities of condensates can mediate intercellular signaling, the BL21 (DE3) competent *E. coli.* was transformed with RLP_SKGP-WT_ plasmid and the DH5α competent *E. coli.* (NEB) was transformed with the DmpR gene circuit. A single colony of the RLP_SKGP-WT_-harboring BL21 (DE3) cells and a single colony of the DmpR-harboring DH5α cells were picked and each grown separately in 2 mL 2× YT liquid medium supplied with 4 % glucose and appropriate antibiotic at 37 °C overnight. The cell culture was back diluted at a ratio of 1:100 with fresh 2× YT liquid medium with appropriate antibiotic. After a 1.5-hour incubation, the BL21 (DE3) cells carrying the RLP_SKGP-WT_ plasmid were either induced or not induced with 0.5 mM IPTG. At 3 h post induction for BL21 cell culture (4.5 h post dilution for 5-alpha cell culture), each cell culture was measured for OD_600_ and adjusted to the same cellular density using M9 Minimal Salts liquid medium (Sigma-Aldrich) supplied with 0.4 % (w/v) sodium acetate, 0.01 % (w/v) thiamine, 2 mM MgSO_4_ and 0.1 mM CaCl_2_ (this medium will be referred as M9 medium in the following content). Each sample was prepared by mixing two cell populations at a volume ratio of 1:2, 1:1, 2:1 with the same total volume. eGFP fluorescence, absorbances at 600 nm and 410 nm were monitored with a 5-min kinetic interval at 37°C for 500 cycles on a TECAN Spark plate reader.

To further confirm the results obtained from plate reader-based monitoring of fluorescence change, we employed confocal microscopy to analyze the fluorescence from cells. Experimentally, we co-cultured BL21 (DE3) competent *E. coli* containing RLP_SKGP-WT_ plasmid with or without condensates with DH5α competent *E. coli.* (NEB) transformed with the DmpR gene circuit, in the M9 medium with pNPP (4 mM). The co-culturing process followed the aforementioned protocol and the initial population ratio between two types of cells was 1:1. After co-culturing, the cells were diluted and subjected to confocal microscopy for further characterization. The fluorescence of EGFP was detected and the images were analyzed with ImageJ (NIH).

## References

1. Shin, Y., and Brangwynne, C.P. (2017). Liquid phase condensation in cell physiology and disease. Science 357, eaaf4382. 10.1126/science.aaf4382.

2. Yang, P., Mathieu, C., Kolaitis, R.-M., Zhang, P., Messing, J., Yurtsever, U., Yang, Z., Wu, J., Li, Y., and Pan, Q. (2020). G3BP1 is a tunable switch that triggers phase separation to assemble stress granules. Cell 181, 325–345. e328.

3. Lyon, A.S., Peeples, W.B., and Rosen, M.K. (2021). A framework for understanding the functions of biomolecular condensates across scales. Nature Reviews Molecular Cell Biology 22, 215–235. 10.1038/s41580-020-00303-z.

4. Roden, C., and Gladfelter, A.S. (2021). RNA contributions to the form and function of biomolecular condensates. Nature Reviews Molecular Cell Biology 22, 183–195. 10.1038/s41580-020-0264-6.

5. Wright, P.E., and Dyson, H.J. (2015). Intrinsically disordered proteins in cellular signalling and regulation. Nature Reviews Molecular Cell Biology 16, 18–29. 10.1038/nrm3920.

6. Sabari, B.R., Hyman, A.A., and Hnisz, D. (2024). Functional specificity in biomolecular condensates revealed by genetic complementation. Nature Reviews Genetics. 10.1038/s41576-024-00780-4.

7. Lyons, H., Veettil, R.T., Pradhan, P., Fornero, C., De La Cruz, N., Ito, K., Eppert, M., Roeder, R.G., and Sabari, B.R. (2023). Functional partitioning of transcriptional regulators by patterned charge blocks. Cell 186, 327–345.e328. 10.1016/j.cell.2022.12.013.

8. Li, P., Banjade, S., Cheng, H.C., Kim, S., Chen, B., Guo, L., Llaguno, M., Hollingsworth, J.V., King, D.S., Banani, S.F., et al. (2012). Phase transitions in the assembly of multivalent signalling proteins. Nature 483, 336–340. 10.1038/nature10879.

9. Harmon, T.S., Holehouse, A.S., Rosen, M.K., and Pappu, R.V. (2017). Intrinsically disordered linkers determine the interplay between phase separation and gelation in multivalent proteins. eLife 6, 30294. 10.7554/eLife.30294.

10. Pappu, R.V., Cohen, S.R., Dar, F., Farag, M., and Kar, M. (2023). Phase Transitions of Associative Biomacromolecules. Chemical Reviews 123, 8945–8987. 10.1021/acs.chemrev.2c00814.

11. King, M.R., Ruff, K.M., Lin, A.Z., Pant, A., Farag, M., Lalmansingh, J.M., Wu, T., Fossat, M.J., Ouyang, W., Lew, M.D., et al. (2024). Macromolecular Condensation Organizes Nucleolar Sub-Phases to Set Up a pH Gradient. Cell 187, 1889–1906. 10.1016/j.cell.2024.02.029.

12. King, M.R., Ruff, K.M., and Pappu, R.V. (2024). Emergent microenvironments of nucleoli. Nucleus 15, 2319957. 10.1080/19491034.2024.2319957.

13. Ruff, K.M., King, M.R., Ying, A.W., Liu, V., Pant, A., Lieberman, W.E., Shinn, M.K., Su, X., Kadoch, C., and Pappu, R.V. (2025). Molecular grammars of intrinsically disordered regions that span the human proteome. bioRxiv, 2025.2002.2027.640591. 10.1101/2025.02.27.640591.

14. Dai, Y., Chamberlayne, C.F., Messina, M.S., Chang, C.J., Zare, R.N., You, L., and Chilkoti, A. (2023). Interface of biomolecular condensates modulates redox reactions. Chem 9, 1594–1609. 10.1016/j.chempr.2023.04.001.

15. Wu, T., King, M.R., Farag, M., Pappu, R.V., and Lew, M.D. (2023). Single fluorogen imaging reveals spatial inhomogeneities within biomolecular condensates. bioRxiv, 2023.2001.2026.525727. 10.1101/2023.01.26.525727.

16. Cakmak, F.P., Choi, S., Meyer, M.O., Bevilacqua, P.C., and Keating, C.D. (2020). Prebiotically-relevant low polyion multivalency can improve functionality of membraneless compartments. Nature Communications 11, 5949. 10.1038/s41467-020-19775-w.

17. Ye, S., Latham, A.P., Tang, Y., Hsiung, C.-H., Chen, J., Luo, F., Liu, Y., Zhang, B., and Zhang, X. (2023). Micropolarity governs the structural organization of biomolecular condensates. bioRxiv, 2023.2003.2030.534881. 10.1101/2023.03.30.534881.

18. Ausserwöger, H., Qian, D., Krainer, G., Csilléry, E.d., Welsh, T.J., Sneideris, T., Franzmann, T.M., Qamar, S., Erkamp, N.A., Nixon-Abell, J., et al. (2023). Quantifying collective interactions in biomolecular phase separation. bioRxiv, 2023.2005.2031.543137. 10.1101/2023.05.31.543137.

19. Klein, I.A., Boija, A., Afeyan, L.K., Hawken, S.W., Fan, M., Dall’Agnese, A., Oksuz, O., Henninger, J.E., Shrinivas, K., Sabari, B.R., et al. (2020). Partitioning of cancer therapeutics in nuclear condensates. Science 368, 1386–1392. 10.1126/science.aaz4427.

20. Kilgore, H.R., and Young, R.A. (2022). Learning the chemical grammar of biomolecular condensates. Nature Chemical Biology. 10.1038/s41589-022-01046-y.

21. Kilgore, H.R., Mikhael, P.G., Overholt, K.J., Boija, A., Hannett, N.M., Van Dongen, C., Lee, T.I., Chang, Y.-T., Barzilay, R., and Young, R.A. (2024). Distinct chemical environments in biomolecular condensates. Nature Chemical Biology 20, 291–301.

22. Posey, A.E., Bremer, A., Erkamp, N.A., Pant, A., Knowles, T., Dai, Y., Mittag, T., and Pappu, R.V. (2024). Biomolecular condensates are characterized by interphase electric potentials. Journal of the American Chemical Society 146, 28268–28281. 10.1021/jacs.4c08946.

23. Dai, Y., Wang, Z.-G., and Zare, R.N. (2024). Unlocking the electrochemical functions of biomolecular condensates. Nature Chemical Biology. 10.1038/s41589-024-01717-y.

24. Adair, G.S. (1923). On the Donnan Equilibrium and the Equation of Gibbs. Science 58, 13–13. doi:10.1126/science.58.1488.13.a.

25. Donnan, F.G. (1924). The Theory of Membrane Equilibria. Chemical Reviews 1, 73–90. 10.1021/cr60001a003.

26. Aydogan Gokturk, P., Sujanani, R., Qian, J., Wang, Y., Katz, L.E., Freeman, B.D., and Crumlin, E.J. (2022). The Donnan potential revealed. Nature Communications 13, 5880. 10.1038/s41467-022-33592-3.

27. Hoffmann, C., Ruff, K.M., Edu, I.A., Shinn, M.K., Tromm, J.V., King, M.R., Pant, A., Ausserwöger, H., Morgan, J.R., Knowles, T.P.J., et al. (2025). Synapsin condensation is governed by sequence-encoded molecular grammars. Journal of Molecular Biology 437, 168987. 10.1016/j.jmb.2025.168987.

28. Farag, M., Cohen, S.R., Borcherds, W.M., Bremer, A., Mittag, T., and Pappu, R.V. (2022). Condensates formed by prion-like low-complexity domains have small-world network structures and interfaces defined by expanded conformations. Nature Communications 13, 7722. 10.1038/s41467-022-35370-7.

29. Welsh, T.J., Krainer, G., Espinosa, J.R., Joseph, J.A., Sridhar, A., Jahnel, M., Arter, W.E., Saar, K.L., Alberti, S., Collepardo-Guevara, R., and Knowles, T.P.J. (2022). Surface Electrostatics Govern the Emulsion Stability of Biomolecular Condensates. Nano Letters 22, 612–621. 10.1021/acs.nanolett.1c03138.

30. Hoffmann, C., Murastov, G., Tromm, J.V., Moog, J.-B., Aslam, M.A., Matkovic, A., and Milovanovic, D. (2023). Electric Potential at the Interface of Membraneless Organelles Gauged by Graphene. Nano Letters 23, 10796–10801. 10.1021/acs.nanolett.3c02915.

31. Chen, M.W., Ren, X., Song, X., Qian, N., Ma, Y., Yu, W., Yang, L., Min, W., Zare, R.N., and Dai, Y. (2025). Transition-State-Dependent Spontaneous Generation of Reactive Oxygen Species by Aβ Assemblies Encodes a Self-Regulated Positive Feedback Loop for Aggregate Formation. Journal of the American Chemical Society. 10.1021/jacs.4c15532.

32. Stroberg, W., and Schnell, S. (2018). Do Cellular Condensates Accelerate Biochemical Reactions? Lessons from Microdroplet Chemistry. Biophysical Journal 115, 3–8. 10.1016/j.bpj.2018.05.023.

33. Mehrgardi, M.A., Mofidfar, M., and Zare, R.N. (2022). Sprayed Water Microdroplets Are Able to Generate Hydrogen Peroxide Spontaneously. Journal of the American Chemical Society 144, 7606–7609. 10.1021/jacs.2c02890.

34. Lee, J.K., Walker, K.L., Han, H.S., Kang, J., Prinz, F.B., Waymouth, R.M., Nam, H.G., and Zare, R.N. (2019). Spontaneous generation of hydrogen peroxide from aqueous microdroplets. Proceedings of the National Academy of Sciences 116, 19294–19298. doi:10.1073/pnas.1911883116.

35. Lee, J.K., Samanta, D., Nam, H.G., and Zare, R.N. (2019). Micrometer-Sized Water Droplets Induce Spontaneous Reduction. Journal of the American Chemical Society 141, 10585–10589. 10.1021/jacs.9b03227.

36. Lewis, L.N. (1993). Chemical catalysis by colloids and clusters. Chemical Reviews 93, 2693–2730.

37. Banani, S.F., Lee, H.O., Hyman, A.A., and Rosen, M.K. (2017). Biomolecular condensates: organizers of cellular biochemistry. Nature Reviews in Molecular and Cell Biology 18, 285–298. 10.1038/nrm.2017.7.

38. Alberti, S., and Hyman, A.A. (2021). Biomolecular condensates at the nexus of cellular stress, protein aggregation disease and ageing. Nature Reviews Molecular Cell Biology 22, 196–213. 10.1038/s41580-020-00326-6.

39. Warshel, A. (1981). Electrostatic basis of structure-function correlation in proteins. Accounts of chemical research 14, 284–290.

40. Warshel, A., Sharma, P.K., Kato, M., Xiang, Y., Liu, H., and Olsson, M.H.M. (2006). Electrostatic Basis for Enzyme Catalysis. Chemical Reviews 106, 3210–3235. 10.1021/cr0503106.

41. Kahraman, A., Morris, R.J., Laskowski, R.A., Favia, A.D., and Thornton, J.M. (2010). On the diversity of physicochemical environments experienced by identical ligands in binding pockets of unrelated proteins. Proteins: Structure, Function, and Bioinformatics 78, 1120–1136.

42. Fried, S.D., and Boxer, S.G. (2017). Electric Fields and Enzyme Catalysis. Annual Review of Biochemistry 86, 387–415. 10.1146/annurev-biochem-061516-044432.

43. Dai, Y., Farag, M., Lee, D., Zeng, X., Kim, K., Son, H.-i., Guo, X., Su, J., Peterson, N., Mohammed, J., et al. (2023). Programmable synthetic biomolecular condensates for cellular control. Nature Chemical Biology 19, 518–528. 10.1038/s41589-022-01252-8.

44. Dzuricky, M., Rogers, B.A., Shahid, A., Cremer, P.S., and Chilkoti, A. (2020). De novo engineering of intracellular condensates using artificial disordered proteins. Nature Chemistry 12, 814–825. 10.1038/s41557-020-0511-7.

45. Zeng, X., Liu, C., Fossat, M.J., Ren, P., Chilkoti, A., and Pappu, R.V. (2021). Design of intrinsically disordered proteins that undergo phase transitions with lower critical solution temperatures. APL Materials 9, 021119. 10.1063/5.0037438.

46. Ruff, K.M., Roberts, S., Chilkoti, A., and Pappu, R.V. (2018). Advances in Understanding Stimulus-Responsive Phase Behavior of Intrinsically Disordered Protein Polymers. Journal of Molecular Biology 430, 4619–4635. 10.1016/j.jmb.2018.06.031.

47. Dai, Y., You, L., and Chilkoti, A. (2023). Engineering synthetic biomolecular condensates. Nature Reviews Bioengineering 1, 466–480. 10.1038/s44222-023-00052-6.

48. McNaught, A.D., and Wilkinson, A. (1997). Compendium of chemical terminology (Blackwell Science Oxford).

49. Cordes, E., and Bull, H. (1974). Mechanism and catalysis for hydrolysis of acetals, ketals, and ortho esters. Chemical Reviews 74, 581–603.

50. Vincent, J.B., Crowder, M.W., and Averill, B.A. (1992). Hydrolysis of phosphate monoesters: a biological problem with multiple chemical solutions. Trends in Biochemical Sciences 17, 105–110. 10.1016/0968-0004(92)90246-6.

51. Lipkin, D., Talbert, P.T., and Cohn, M. (1954). The mechanism of the alkaline hydrolysis of ribonucleic acids. Journal of the American Chemical Society 76, 2871–2872.

52. Zastrow, M.L., Peacock, A.F., Stuckey, J.A., and Pecoraro, V.L. (2012). Hydrolytic catalysis and structural stabilization in a designed metalloprotein. Nature chemistry 4, 118–123.

53. Twigg, M.V. (2018). Catalyst handbook (Routledge).

54. Zwicker, D., Seyboldt, R., Weber, C.A., Hyman, A.A., and Jülicher, F. (2017). Growth and division of active droplets provides a model for protocells. Nature Physics 13, 408–413. 10.1038/nphys3984.

55. Wang, Z., Danovich, D., Ramanan, R., and Shaik, S. (2018). Oriented-External Electric Fields Create Absolute Enantioselectivity in Diels–Alder Reactions: Importance of the Molecular Dipole Moment. Journal of the American Chemical Society 140, 13350–13359. 10.1021/jacs.8b08233.

56. Shaik, S., Mandal, D., and Ramanan, R. (2016). Oriented electric fields as future smart reagents in chemistry. Nature Chemistry 8, 1091–1098. 10.1038/nchem.2651.

57. Epifanovsky, E., Gilbert, A.T., Feng, X., Lee, J., Mao, Y., Mardirossian, N., Pokhilko, P., White, A.F., Coons, M.P., and Dempwolff, A.L. (2021). Software for the frontiers of quantum chemistry: An overview of developments in the Q-Chem 5 package. The Journal of chemical physics 155.

58. Fried, S.D., and Boxer, S.G. (2015). Measuring Electric Fields and Noncovalent Interactions Using the Vibrational Stark Effect. Accounts of Chemical Research 48, 998–1006. 10.1021/ar500464j.

59. Zuo, Y.-X., and Stenby, E.H. (1996). A Linear Gradient Theory Model for Calculating Interfacial Tensions of Mixtures. Journal of Colloid and Interface Science 182, 126–132. 10.1006/jcis.1996.0443.

60. Bastos-González, D., Pérez-Fuentes, L., Drummond, C., and Faraudo, J. (2016). Ions at interfaces: the central role of hydration and hydrophobicity. Current Opinion in Colloid & Interface Science 23, 19–28. 10.1016/j.cocis.2016.05.010.

61. Lyklema, J. (2005). Fundamentals of interface and colloid science: soft colloids (Elsevier).

62. Parsons, R. (1990). The electrical double layer: recent experimental and theoretical developments. Chemical Reviews 90, 813–826.

63. Henderson, D., and Boda, D. (2009). Insights from theory and simulation on the electrical double layer. Physical chemistry chemical physics 11, 3822–3830.

64. Berg, J.C. (2010). An introduction to interfaces & colloids: the bridge to nanoscience (World Scientific).

65. Choi, J.-M., Dar, F., and Pappu, R.V. (2019). LASSI: A lattice model for simulating phase transitions of multivalent proteins. PLoS computational biology 15, e1007028.

66. Farag, M., Borcherds, W.M., Bremer, A., Mittag, T., and Pappu, R.V. (2023). Phase separation of protein mixtures is driven by the interplay of homotypic and heterotypic interactions. Nature Communications 14, 5527. 10.1038/s41467-023-41274-x.

67. Martin, E.W., Holehouse, A.S., Peran, I., Farag, M., Incicco, J.J., Bremer, A., Grace, C.R., Soranno, A., Pappu, R.V., and Mittag, T. (2020). Valence and patterning of aromatic residues determine the phase behavior of prion-like domains. Science 367, 694–699. 10.1126/science.aaw8653.

68. Bremer, A., Farag, M., Borcherds, W.M., Peran, I., Martin, E.W., Pappu, R.V., and Mittag, T. (2022). Deciphering how naturally occurring sequence features impact the phase behaviours of disordered prion-like domains. Nature Chemistry 14, 196–207. 10.1038/s41557-021-00840-w.

69. Leunissen, M.E., Blaaderen, A.v., Hollingsworth, A.D., Sullivan, M.T., and Chaikin, P.M. (2007). Electrostatics at the oil-water interface, stability, and order in emulsions and colloids. Proceedings of the National Academy of Sciences 104, 2585–2590. doi:10.1073/pnas.0610589104.

70. Spohr, E. (1997). Effect of electrostatic boundary conditions and system size on the interfacial properties of water and aqueous solutions. The Journal of chemical physics 107, 6342–6348.

71. Hortigon-Vinagre, M.P., Zamora, V., Burton, F.L., Green, J., Gintant, G.A., and Smith, G.L. (2016). The Use of Ratiometric Fluorescence Measurements of the Voltage Sensitive Dye Di-4-ANEPPS to Examine Action Potential Characteristics and Drug Effects on Human Induced Pluripotent Stem Cell-Derived Cardiomyocytes. Toxicological Sciences 154, 320–331. 10.1093/toxsci/kfw171.

72. Tominaga, T., Tominaga, Y., Yamada, H., Matsumoto, G., and Ichikawa, M. (2000). Quantification of optical signals with electrophysiological signals in neural activities of Di-4-ANEPPS stained rat hippocampal slices. Journal of neuroscience methods 102, 11–23.

73. Kar, M., Dar, F., Welsh, T.J., Vogel, L.T., Kühnemuth, R., Majumdar, A., Krainer, G., Franzmann, T.M., Alberti, S., Seidel, C.A.M., et al. (2022). Phase-separating RNA-binding proteins form heterogeneous distributions of clusters in subsaturated solutions. Proceedings of the National Academy of Sciences 119, e2202222119. doi:10.1073/pnas.2202222119.

74. Li, Z., Shen, Q., Usher, E.T., Anderson, A.P., Iburg, M., Lin, R., Zimmer, B., Meyer, M.D., Holehouse, A.S., You, L., et al. (2024). Phase transition of GvpU regulates gas vesicle clustering in bacteria. Nature Microbiology 9, 1021–1035. 10.1038/s41564-024-01648-3.

75. Yanas, A., Shweta, H., Owens, M.C., Liu, K.F., and Goldman, Y.E. (2024). RNA helicases DDX3X and DDX3Y form nanometer-scale RNA-protein clusters that support catalytic activity. Current Biology 34, 5714–5727.e5716. 10.1016/j.cub.2024.10.055.

76. Jungwirth, P., and Winter, B. (2008). Ions at Aqueous Interfaces: From Water Surface to Hydrated Proteins. Annual Review of Physical Chemistry 59, 343–366. 10.1146/annurev.physchem.59.032607.093749.

77. Ng, S., Howshall, C., Ho, T.N., Mai, B.K., Zhou, Y., Qin, C., Tee, K.Z., Liu, P., Romiti, F., and Hoveyda, A.H. (2024). Catalytic prenyl conjugate additions for synthesis of enantiomerically enriched PPAPs. Science 386, 167–175. doi:10.1126/science.adr8612.

78. Butré, C.I., Sforza, S., Gruppen, H., and Wierenga, P.A. (2014). Introducing enzyme selectivity: a quantitative parameter to describe enzymatic protein hydrolysis. Analytical and Bioanalytical Chemistry 406, 5827–5841. 10.1007/s00216-014-8006-2.

79. Oldham, K.B. (2008). A Gouy–Chapman–Stern model of the double layer at a (metal)/(ionic liquid) interface. Journal of Electroanalytical Chemistry 613, 131–138.

80. Onsager, L. (1936). Electric Moments of Molecules in Liquids. Journal of the American Chemical Society 58, 1486–1493. 10.1021/ja01299a050.

81. Sorenson, S.A., Patrow, J.G., and Dawlaty, J.M. (2017). Solvation Reaction Field at the Interface Measured by Vibrational Sum Frequency Generation Spectroscopy. Journal of the American Chemical Society 139, 2369–2378. 10.1021/jacs.6b11940.

82. Shi, L., Hu, F., and Min, W. (2019). Optical mapping of biological water in single live cells by stimulated Raman excited fluorescence microscopy. Nature communications 10, 4764.

83. Xiong, H., Lee, J.K., Zare, R.N., and Min, W. (2020). Strong Electric Field Observed at the Interface of Aqueous Microdroplets. The Journal of Physical Chemistry Letters 11, 7423–7428. 10.1021/acs.jpclett.0c02061.

84. Wei, L., Yu, Y., Shen, Y., Wang, M.C., and Min, W. (2013). Vibrational imaging of newly synthesized proteins in live cells by stimulated Raman scattering microscopy. Proceedings of the National Academy of Sciences 110, 11226–11231.

85. Xiong, H., Shi, L., Wei, L., Shen, Y., Long, R., Zhao, Z., and Min, W. (2019). Stimulated Raman excited fluorescence spectroscopy and imaging. Nature photonics 13, 412–417.

86. Wei, L., Hu, F., Chen, Z., Shen, Y., Zhang, L., and Min, W. (2016). Live-cell bioorthogonal chemical imaging: stimulated Raman scattering microscopy of vibrational probes. Accounts of chemical research 49, 1494–1502.

87. Freudiger, C.W., Min, W., Saar, B.G., Lu, S., Holtom, G.R., He, C., Tsai, J.C., Kang, J.X., and Xie, X.S. (2008). Label-free biomedical imaging with high sensitivity by stimulated Raman scattering microscopy. Science 322, 1857–1861.

88. Min, W., and Gao, X. (2024). The Duality of Raman Scattering. Accounts of Chemical Research 57, 1896–1905.

89. Lang, X., Shi, L., Zhao, Z., and Min, W. (2024). Probing the structure of water in individual living cells. Nature Communications 15, 5271. 10.1038/s41467-024-49404-9.

90. Shi, L., LaCour, R.A., Lang, X., Heindel, J.P., Head-Gordon, T., and Min, W. (2024). Water Structure and Electric Fields at the Interface of Oil Droplets. arXiv preprint arXiv:2405.02207.

91. Hua, M., Peng, Z., Guha, R.D., Ruan, X., Ng, K.C., Demarteau, J., Haber, S., Fricke, S.N., Reimer, J.A., Salmeron, M.B., et al. (2024). Mechanochemically accelerated deconstruction of chemically recyclable plastics. Science Advances 10, eadq3801. doi:10.1126/sciadv.adq3801.

92. Henao, A., Ruiz, G.N., Steinke, N., Cerveny, S., Macovez, R., Guàrdia, E., Busch, S., McLain, S.E., Lorenz, C.D., and Pardo, L.C. (2020). On the microscopic origin of the cryoprotective effect in lysine solutions. Physical Chemistry Chemical Physics 22, 6919–6927. 10.1039/C9CP06192D.

93. Nott, T.J., Petsalaki, E., Farber, P., Jervis, D., Fussner, E., Plochowietz, A., Craggs, T.D., Bazett-Jones, D.P., Pawson, T., and Forman-Kay, J.D. (2015). Phase transition of a disordered nuage protein generates environmentally responsive membraneless organelles. Molecular cell 57, 936–947.

94. Bashkin, J.K. (1999). Hydrolysis of phosphates, esters and related substrates by models of biological catalysts. Current Opinion in Chemical Biology 3, 752–758. 10.1016/S1367-5931(99)00036-8.

95. Morth, J.P., Pedersen, B.P., Buch-Pedersen, M.J., Andersen, J.P., Vilsen, B., Palmgren, M.G., and Nissen, P. (2011). A structural overview of the plasma membrane Na+, K+-ATPase and H+-ATPase ion pumps. Nature reviews Molecular cell biology 12, 60–70.

96. Kühlbrandt, W. (2004). Biology, structure and mechanism of P-type ATPases. Nature reviews Molecular cell biology 5, 282–295.

97. Saveant, J.M. (1993). Electron transfer, bond breaking, and bond formation. Accounts of chemical research 26, 455–461.

98. Dai, Y., Zhou, Z., Yu, W., Ma, Y., Kim, K., Rivera, N., Mohammed, J., Lantelme, E., Hsu-Kim, H., Chilkoti, A., and You, L. (2024). Biomolecular condensates regulate cellular electrochemical equilibria. Cell 187, 5951–5966.e5918. 10.1016/j.cell.2024.08.018.

99. Weber, J., Li, Z., and Rinas, U. (2021). Recombinant protein production provoked accumulation of ATP, fructose-1,6-bisphosphate and pyruvate in E. coli K12 strain TG1. Microbial Cell Factories 20, 169. 10.1186/s12934-021-01661-9.

100. Bhattacharya, S.K., and Dubey, A.K. (1995). Metabolic burden as reflected by maintenance coefficient of recombinant Escherichia coli overexpressing target gene. Biotechnology letters 17, 1155–1160.

101. Wu, G., Yan, Q., Jones, J.A., Tang, Y.J., Fong, S.S., and Koffas, M.A. (2016). Metabolic burden: cornerstones in synthetic biology and metabolic engineering applications. Trends in biotechnology 34, 652–664.

102. Groisman, E.A. (2001). The pleiotropic two-component regulatory system PhoP-PhoQ. Journal of bacteriology 183, 1835–1842.

103. Liu, W., and Hulett, F.M. (1997). Bacillus subtilis PhoP binds to the phoB tandem promoter exclusively within the phosphate starvation-inducible promoter. Journal of bacteriology 179, 6302–6310.

104. Makino, K., Shinagawa, H., Amemura, M., Kimura, S., Nakata, A., and Ishihama, A. (1988). Regulation of the phosphate regulon of Escherichia coli: activation of pstS transcription by PhoB protein in vitro. Journal of molecular biology 203, 85–95.

105. Wanner, B. (1993). Gene regulation by phosphate in enteric bacteria. Journal of cellular biochemistry 51, 47–54.

106. Baek, J.-H., and Lee, S.-Y. (2007). Transcriptome analysis of phosphate starvation response in Escherichia coli. Journal of microbiology and biotechnology 17, 244–252.

107. Yang, C., Huang, T.-W., Wen, S.-Y., Chang, C.-Y., Tsai, S.-F., Wu, W.-F., and Chang, C.- H. (2012). Genome-wide PhoB binding and gene expression profiles reveal the hierarchical gene regulatory network of phosphate starvation in Escherichia coli.

108. Zwir, I., Shin, D., Kato, A., Nishino, K., Latifi, T., Solomon, F., Hare, J.M., Huang, H., and Groisman, E.A. (2005). Dissecting the PhoP regulatory network of Escherichia coli and Salmonella enterica. Proceedings of the National Academy of Sciences 102, 2862–2867.

109. Liu, J., Prindle, A., Humphries, J., Gabalda-Sagarra, M., Asally, M., Lee, D.-y.D., Ly, S., Garcia-Ojalvo, J., and Süel, G.M. (2015). Metabolic co-dependence gives rise to collective oscillations within biofilms. Nature 523, 550–554. 10.1038/nature14660.

110. Humphries, J., Xiong, L., Liu, J., Prindle, A., Yuan, F., Arjes, H.A., Tsimring, L., and Süel, G.M. (2017). Species-Independent Attraction to Biofilms through Electrical Signaling. Cell 168, 200–209.e212. 10.1016/j.cell.2016.12.014.

111. Andrianantoandro, E., Basu, S., Karig, D.K., and Weiss, R. (2006). Synthetic biology: new engineering rules for an emerging discipline. Molecular systems biology 2, 2006.0028.

112. Cao, Y., Lopatkin, A., and You, L. (2016). Elements of biological oscillations in time and space. Nature Structural & Molecular Biology 23, 1030–1034.

113. Cao, Y., Ryser, Marc D., Payne, S., Li, B., Rao, Christopher V., and You, L. (2016). Collective Space-Sensing Coordinates Pattern Scaling in Engineered Bacteria. Cell 165, 620–630. 10.1016/j.cell.2016.03.006.

114. Sun, S., Peng, K., Sun, S., Wang, M., Shao, Y., Li, L., Xiang, J., Sedjoah, R.-C.A.-A., and Xin, Z. (2023). Engineering Modular and Highly Sensitive Cell-Based Biosensors for Aromatic Contaminant Monitoring and High-Throughput Enzyme Screening. ACS Synthetic Biology 12, 877–891. 10.1021/acssynbio.3c00036.

115. Sarand, I., Skärfstad, E., Forsman, M., Romantschuk, M., and Shingler, V. (2001). Role of the DmpR-Mediated Regulatory Circuit in Bacterial Biodegradation Properties in Methylphenol-Amended Soils. Applied and Environmental Microbiology 67, 162–171. doi:10.1128/AEM.67.1.162-171.2001.

116. Putnam, A., Thomas, L., and Seydoux, G. (2023). RNA granules: functional compartments or incidental condensates? Genes & Development 37, 354–376. 10.1101/gad.350518.123.

117. Ausserwöger, H., Qian, D., Krainer, G., Welsh, T.J., Sneideris, T., Franzmann, T.M., Qamar, S., Nixon-Abell, J., Kar, M., George-Hyslop, P.S., et al. (2023). Condensate partitioning governs the mechanism of action of FUS phase separation modulators. bioRxiv, 2023.2005.2031.543137. 10.1101/2023.05.31.543137.

118. Van Hemmen, J., and Bleichrodt, J. (1971). The decomposition of adenine by ionizing radiation. Radiation Research 46, 444–456.

119. Yan, X., Kuster, D., Mohanty, P., Nijssen, J., Pombo-García, K., Rizuan, A., Franzmann, T.M., Sergeeva, A., Passos, P.M., George, L., et al. (2024). Intra-condensate demixing of TDP-43 inside stress granules generates pathological aggregates. bioRxiv, 2024.2001.2023.576837. 10.1101/2024.01.23.576837.

120. Khalil, A.S., and Collins, J.J. (2010). Synthetic biology: applications come of age. Nature Reviews Genetics 11, 367–379. 10.1038/nrg2775.

121. Wessén, J., Pal, T., and Chan, H.S. (2022). Field theory description of ion association in re-entrant phase separation of polyampholytes. The Journal of Chemical Physics 156.

122. Friedowitz, S., Lou, J., Barker, K.P., Will, K., Xia, Y., and Qin, J. (2021). Looping-in complexation and ion partitioning in nonstoichiometric polyelectrolyte mixtures. Science Advances 7, eabg8654. doi:10.1126/sciadv.abg8654.

123. Schlenoff, J.B., Yang, M., Digby, Z.A., and Wang, Q. (2019). Ion Content of Polyelectrolyte Complex Coacervates and the Donnan Equilibrium. Macromolecules 52, 9149–9159. 10.1021/acs.macromol.9b01755.

124. Neitzel, A.E., Fang, Y.N., Yu, B., Rumyantsev, A.M., De Pablo, J.J., and Tirrell, M.V. (2021). Polyelectrolyte complex coacervation across a broad range of charge densities. Macromolecules 54, 6878–6890.

125. Ghasemi, M., Friedowitz, S., and Larson, R.G. (2020). Analysis of partitioning of salt through doping of polyelectrolyte complex coacervates. Macromolecules 53, 6928–6945.

126. Li, L., Srivastava, S., Andreev, M., Marciel, A.B., de Pablo, J.J., and Tirrell, M.V. (2018). Phase Behavior and Salt Partitioning in Polyelectrolyte Complex Coacervates. Macromolecules 51, 2988–2995. 10.1021/acs.macromol.8b00238.

127. Zhang, P., Alsaifi, N.M., Wu, J., and Wang, Z.-G. (2016). Salting-Out and Salting-In of Polyelectrolyte Solutions: A Liquid-State Theory Study. Macromolecules 49, 9720–9730. 10.1021/acs.macromol.6b02160.

128. Zhang, P., Shen, K., Alsaifi, N.M., and Wang, Z.-G. (2018). Salt partitioning in complex coacervation of symmetric polyelectrolytes. Macromolecules 51, 5586–5593.

129. Zhang, P., Alsaifi, N.M., Wu, J., and Wang, Z.-G. (2018). Polyelectrolyte complex coacervation: Effects of concentration asymmetry. The Journal of Chemical Physics 149. 10.1063/1.5028524.

130. Holehouse, A.S., and Kragelund, B.B. (2024). The molecular basis for cellular function of intrinsically disordered protein regions. Nature Reviews Molecular Cell Biology 25, 187–211. 10.1038/s41580-023-00673-0.

131. Nott, T.J., Craggs, T.D., and Baldwin, A.J. (2016). Membraneless organelles can melt nucleic acid duplexes and act as biomolecular filters. Nature Chemistry 8, 569–575. 10.1038/nchem.2519.

132. Abbas, M., Lipiński, W.P., Nakashima, K.K., Huck, W.T.S., and Spruijt, E. (2021). A short peptide synthon for liquid–liquid phase separation. Nature Chemistry 13, 1046–1054. 10.1038/s41557-021-00788-x.

133. Zepik, H.H., Blöchliger, E., and Luisi, P.L. (2001). A chemical model of homeostasis. Angewandte Chemie International Edition 40, 199–202.

134. Pittendrigh, C.S., and Caldarola, P.C. (1973). General homeostasis of the frequency of circadian oscillations. Proceedings of the National Academy of Sciences 70, 2697–2701.

135. Guggenheim, E. (1940). The thermodynamics of interfaces in systems of several components. Transactions of the Faraday Society 35, 397–412.

136. Hansen, R.S. (1962). Thermodynamics OF interfaces between condensed PHASES1. The Journal of Physical Chemistry 66, 410–415.

137. Zhang, P., and Wang, Z.-G. (2021). Interfacial Structure and Tension of Polyelectrolyte Complex Coacervates. Macromolecules 54, 10994–11007. 10.1021/acs.macromol.1c01809.

138. Vis, M., Peters, V.F., Tromp, R.H., and Erné, B.H. (2014). Donnan potentials in aqueous phase-separated polymer mixtures. Langmuir 30, 5755–5762.

139. Hummer, G., Pratt, L.R., and Garcia, A.E. (1996). Free energy of ionic hydration. The Journal of Physical Chemistry 100, 1206–1215.

140. Carnie, S.L., and Torrie, G.M. (1984). The statistical mechanics of the electrical double layer. Advances in Chemical Physics, 141–253.

141. Folkmann, A.W., Putnam, A., Lee, C.F., and Seydoux, G. (2021). Regulation of biomolecular condensates by interfacial protein clusters. Science 373, 1218–1224. doi:10.1126/science.abg7071.

142. Grahame, D.C. (1947). The electrical double layer and the theory of electrocapillarity. Chemical reviews 41, 441–501.

143. Arad, E., and Jelinek, R. (2022). Catalytic amyloids. Trends in Chemistry 4, 907–917. 10.1016/j.trechm.2022.07.001.

144. Joshi, A., Avni, A., Walimbe, A., Rai, S.K., Sarkar, S., and Mukhopadhyay, S. (2024). Hydrogen-Bonded Network of Water in Phase-Separated Biomolecular Condensates. The Journal of Physical Chemistry Letters, 7724–7734. 10.1021/acs.jpclett.4c01153.

145. Posey, A.E., Bremer, A., Erkamp, N.A., Pant, A., Knowles, T.P.J., Dai, Y., Mittag, T., and Pappu, R.V. (2024). Biomolecular Condensates are Characterized by Interphase Electric Potentials. Journal of the American Chemical Society. 10.1021/jacs.4c08946.

146. Kahana, A., and Lancet, D. (2021). Self-reproducing catalytic micelles as nanoscopic protocell precursors. Nature Reviews Chemistry 5, 870–878. 10.1038/s41570-021-00329-7.

147. Li, C.-J., and Chen, L. (2006). Organic chemistry in water. Chemical Society Reviews 35, 68–82.

148. Kitanosono, T., and Kobayashi, S. (2020). Reactions in water involving the “On-Water” mechanism. Chemistry–A European Journal 26, 9408–9429.

149. Rufo, C.M., Moroz, Y.S., Moroz, O.V., Stöhr, J., Smith, T.A., Hu, X., DeGrado, W.F., and Korendovych, I.V. (2014). Short peptides self-assemble to produce catalytic amyloids. Nature chemistry 6, 303–309.

150. Makam, P., Yamijala, S.S.R.K.C., Tao, K., Shimon, L.J.W., Eisenberg, D.S., Sawaya, M.R., Wong, B.M., and Gazit, E. (2019). Non-proteinaceous hydrolase comprised of a phenylalanine metallo-supramolecular amyloid-like structure. Nature Catalysis 2, 977–985. 10.1038/s41929-019-0348-x.

151. Zhou, H., Huertas, J., Maristany, M.J., Russell, K., Hwang, J.H., Yao, R.-w., Hutchings, J., Shiozaki, M., Zhao, X., Doolittle, L.K., et al. (2025). Multi-scale structure of chromatin condensates rationalizes phase separation and material properties. bioRxiv, 2025.2001.2017.633609. 10.1101/2025.01.17.633609.

152. Keber, F.C., Nguyen, T., Mariossi, A., Brangwynne, C.P., and Wühr, M. (2024). Evidence for widespread cytoplasmic structuring into mesoscale condensates. Nature Cell Biology 26, 346–352. 10.1038/s41556-024-01363-5.

153. Alshareedah, I., Borcherds, W.M., Cohen, S.R., Singh, A., Posey, A.E., Farag, M., Bremer, A., Strout, G.W., Tomares, D.T., Pappu, R.V., et al. (2024). Sequence-specific interactions determine viscoelasticity and aging dynamics of protein condensates. Nature Physics 20, 1482–1491. 10.1038/s41567-024-02558-1.

154. Alshareedah, I., Singh, A., Yang, S., Ramachandran, V., Quinn, A., Potoyan, D.A., and Banerjee, P.R. (2024). Determinants of viscoelasticity and flow activation energy in biomolecular condensates. Science Advances 10, eadi6539. 10.1126/sciadv.adi6539.

155. Cohen, S.R., Banerjee, P.R., and Pappu, R.V. (2024). Direct computations of viscoelastic moduli of biomolecular condensates. Journal of Chemical Physics 161, 095103. 10.1063/5.0223001.

156. Bergeron-Sandoval, L.P., Kumar, S., Heris, H.K., Chang, C.L.A., Cornell, C.E., Keller, S.L., Francois, P., Hendricks, A.G., Ehrlicher, A.J., Pappu, R.V., and Michnick, S.W. (2021). Endocytic proteins with prion-like domains form viscoelastic condensates that enable membrane remodeling. Proceedings of the National Academy of Sciences USA 118, e2113789118. 10.1073/pnas.2113789118.

157. Feric, M., Vaidya, N., Harmon, T.S., Mitrea, D.M., Zhu, L., Richardson, T.M., Kriwacki, R.W., Pappu, R.V., and Brangwynne, C.P. (2016). Coexisting liquid phases underlie nucleolar subcompartments. Cell 165, 1686–1697.

158. Feric, M., Sarfallah, A., Dar, F., Temiakov, D., Pappu, R.V., and Misteli, T. (2022). Mesoscale structure–function relationships in mitochondrial transcriptional condensates. Proceedings of the National Academy of Sciences 119, e2207303119. doi:10.1073/pnas.2207303119.

159. Gouveia, B., Kim, Y., Shaevitz, J.W., Petry, S., Stone, H.A., and Brangwynne, C.P. (2022). Capillary forces generated by biomolecular condensates. Nature 609, 255–264. 10.1038/s41586-022-05138-6.

160. Boyd, J.G., Loufakis, D., and Lutkenhaus, J.L. (2024). Coupled electro-chemo-viscoelastic constitutive model for a supercapacitor electrode. Journal of Applied Physics 136. 10.1063/5.0209577.

161. Quiroz, F.G., and Chilkoti, A. (2015). Sequence heuristics to encode phase behaviour in intrinsically disordered protein polymers. Nature Materials 14, 1164–1171. 10.1038/nmat4418.

162. Hassouneh, W., Christensen, T., and Chilkoti, A. (2010). Elastin-like polypeptides as a purification tag for recombinant proteins. Current protocols in protein science 61, 6.11. 11–16.11. 16.

163. Xie, D., Zhou, Y., Xu, D., and Guo, H. (2005). Solvent effect on concertedness of the transition state in the hydrolysis of p-nitrophenyl acetate. Organic Letters 7, 2093–2095.

164. Matta, M.S., and Toenjes, A.A. (1985). Solvation effects on the alkaline hydrolysis of some p-nitrophenyl esters. Journal of the American Chemical Society 107, 7591–7596.

165. Klamt, A. (1995). Conductor-like screening model for real solvents: a new approach to the quantitative calculation of solvation phenomena. The Journal of Physical Chemistry 99, 2224–2235.

166. Klamt, A., and Schüürmann, G. (1993). COSMO: a new approach to dielectric screening in solvents with explicit expressions for the screening energy and its gradient. Journal of the Chemical Society, Perkin Transactions 2, 799–805.

167. Amin, M., and Küpper, J. (2020). Variations in proteins dielectric constants. ChemistryOpen 9, 691–694.

